# Systematics of the fleshy-fruited Sonerileae (Melastomataceae)

**DOI:** 10.1101/2024.03.03.582138

**Authors:** J. Peter Quakenbush, Luo Chen, Darin S. Penneys, Todd J. Barkman, Ying Liu, Deepthi Yakandawala, Marie-Claire Veranso-Libalah, Gudrun Kadereit

## Abstract

With approximately 1080 species, Sonerileae is the second largest tribe in the Melastomataceae. Approximately 40% of Sonerileae species belong to fleshy-fruited genera (*Catanthera*, *Heteroblemma*, *Kendrickia*, *Medinilla*, *Pachycentria*, and *Plethiandra*). Relatively few species, especially of the fleshy-fruited taxa, have been sampled for phylogenetic study. Consequently, there is huge uncertainty resulting in many unanswered questions about their evolutionary history, including the monophyly of the largest genus, *Medinilla*. In this study, the phylogeny of the fleshy-fruited Sonerileae was reconstructed using 385 nuclear and 81 plastid protein-coding loci recovered from target capture. Our study revealed that the fleshy fruited Sonerileae are polyphyletic and belong to three lineages. *Kendrickia* is sister to an Afrotropical endemic clade. *Heteroblemma* and *Catanthera* belong to a second clade and are most closely related to some *Phyllagathis* and *Driessenia* species. *Medinilla* forms a third clade, and includes *Pachycentria* and *Plethiandra*. Within *Medinilla*, fifteen clades are identified and characterized. To make *Medinilla* monophyletic, the genus is redefined to include *Pachycentria* and *Plethiandra*. Major lineages identified within *Medinilla* lay the groundwork for an infrageneric classification system. Areas of the phylogenetic tree with high conflict or weak sampling are identified to aid further studies in the tribe.

## INTRODUCTION

With nearly 6000 species, the Melastomataceae are among the 10 most species-rich plant families (Ulloa Ulloa & al., 2022). They are an important ecological component of primarily tropical habitats worldwide and serve as a major evolutionary study system (Goldenberg & al., 2022). The second largest tribe in the family is Sonerileae, consisting of well over 1000 species (Liu & al., 2022; Penneys & al., 2022). Generic delimitation has been notoriously problematic in this tribe, hampering understanding of its diversity. Sonerileae currently includes 45 genera, 39 of which have capsular fruits. However, approximately 40% of Sonerileae species have fleshy fruits. Fleshy-fruited genera include *Medinilla* Gaudich. ex DC., *Pachycentria* Blume, *Plethiandra* Hook.f., *Heteroblemma* (Blume) Cámara-Leret, Ridd.-Num. & Veldkamp, *Catanthera* F.Muell., and *Kendrickia* Hook.f.

*Medinilla* is by far the largest and most widely distributed of these fleshy-fruited genera (Liu & al., 2022). It is the most species-rich genus in Sonerileae and second most species-rich genus in Melastomataceae. Nearly 400 species are found in the wet Paleotropics, from Africa to Polynesia. The exact number of species is unclear. The last comprehensive revision of *Medinilla* dates back to Cogniaux (1891), a time when 75% of the species we recognize today were yet to be described. Subsequent revisions are either outdated or absent, leaving significant gaps in our understanding of the genus, particularly in regions with high diversity like Madagascar and New Guinea (Cámara-Leret & al., 2020). Most species are covered in regional lists and revisions. See Veranso-Libalah & al. (2023) for Africa; Perrier de la Bâthie (1951) for Madagascar and the Comoro Islands; Sasidharan & Sujanapal (2005) for the Western Ghats; Bremer (1988) for Sri Lanka; the Flora of China (2013) and Flora of Thailand (2001) for mainland Southeast Asia (including Hainan and Taiwan); Kartonegoro (2022) for Malesia; Merrill & Perry (1943) for the Solomon Islands; & Smith (1985) for Fiji.

*Medinilla* is also the most poorly defined of the fleshy-fruited genera (Liu & al., 2022; Penneys & al., 2022). Regalado (1990, 1995) and Bodegom & Veldkamp (2001) provided a detailed overview of its complex taxonomic history. Currently, *Medinilla* is understood as a heterogeneous group of terrestrial shrubs, climbers, and epiphytes, often distinguished by isomorphic stamens, weakly produced connective, and anther appendages (two ventral, one dorsal; Regalado, 1995). However, exceptions abound and the genus has been notoriously hard to delimit. Regalado (1995) described the situation as a “taxonomic impasse”. He envisioned *Medinilla* as a central plexus from which small satellite genera have been separated. Kadereit (2005) considered the genus “probably polyphyletic”, as did Veranso-Libalah & al. (2022). The latter study was based on the combined sequences of two nuclear and three plastid markers. Seventeen *Medinilla* species were resolved in two clades, though these were not strongly supported. Until recently, *Medinilla* certainly was polyphyletic because *Pseudodissochaeta* (Kartonegoro & al., 2020; Kartonegoro & al., 2021) and *Myrianthemum* (Chen & al., 2023) were considered synonyms of *Medinilla*. Morphological and molecular evidence has helped demonstrate that these two genera belong to the berry-fruited tribe Dissochaeteae. Other studies support the monophyly of *Medinilla* (Maurin & al., 2021; Zhou & al., 2019; Zhou & al., 2022), but all suffer from limited sampling and a serious lack of crucial taxa (e.g., other fleshy-fruited Sonerileae or the above-mentioned satellite genera). The inclusion of sequences from *Pachycentria* introduces a challenge to the presumed monophyly of *Medinilla*, as indicated by studies conducted by Kartonegoro & al. (2021) and Chen & al. (2023). The results suggest that *Medinilla* is paraphyletic when *Pachycentria* is accepted. To thoroughly test the monophyly of *Medinilla*, there is a clear necessity for comprehensive sampling, not only within *Medinilla* itself, but also across other genera.

No comprehensive infrageneric classification system for *Medinilla* exists to guide the sampling; however, many distinctive morphological groups within *Medinilla* have been recognized. *Carionia* Naudin, *Cephalomedinilla* Merr., *Dactyliota* Blume*, Diplogenea* Lindl., *Erpetina* Naudin, *Hypenanthe* Blume, and *Triplectrum* D.Don ex Wight & Arn. are generic names considered synonymous with *Medinilla*. Several sections have been proposed as well: *M.* sect. *Medinilla* (= *M.* sect. *Campsoplacuntia* Blume (1831), nom. inval. = *M.* sect. *Eu-Medinilla* Bakh.f. (1943), nom. inval.), *M.* sect. *Sarcoplacunita* Blume (1831), *M.* sect. *Apateon* Blume (1849), *M.* sect. *Heteromedinilla* Bakh.f. (1943), *M.* sect. *Septatae* (with three subgroups; Perrier de la Bâthie, 1951), and *M.* sect. *Adhaerentes* (with six subgroups; Perrier de la Bâthie, 1951). Informal species alliances have also been recognized. Veldkamp (1978, 1988) revised species in the *M. myrtiformis*-alliance. Regalado treated 11 and 12 species alliances in his revisions of Bornean (1990) and Philippine (1995) *Medinilla*, respectively. In the most comprehensive classification system to date, Clausing (1999) sorted 215 species into Group 1 (with 13 major alliances excluding *Heteroblemma*) and Group 2 (with four major alliances). Soon after, Bodegom & Veldkamp (2001) identified and revised the pseudo-stipular species of *Medinilla*. Despite these efforts, a significant challenge remains because there is considerable overlap of species between many groups, and numerous species are left unassigned. The complexity of *Medinilla’*s diversity necessitates the identification of major lineages within the genus. This task is critical for gaining a more nuanced understanding of the intricate relationships among species and a prerequisite for any further evolutionary study of the Asian Sonerileae.

*Pachycentria* (including *Pogonanthera* Blume) consists of eight species in Malesia, with two of them being widespread (Clausing, 2000). It is characterized by a small ovary in a strongly constricted, urceolate hypanthium and seeds with comb-shaped testa cells (Clausing, 2000). Ventral anther appendages are generally lacking. The dorsal appendage can be frayed, bifurcated, or tufted (*Pogonanthera*). Baillon (1879) considered *Pachycentria* conspecific with *Medinilla* and maintained *Pogonanthera* with doubt. Clausing (2000) combined *Pachycentria* and *Pogonanthera* and tentatively maintained their distinction from *Medinilla*. Kartonegoro & al. (2021) found support (posterior probability (PP) = 1, Bayesian inference; Bootstrap (BS) = 85, maximum likelihood; BS = 65, parsimony analysis) for two *Pachycentria* species being nested among 13 *Medinilla* species, based on the combination of two nuclear and four chloroplast markers. Using the same markers and a maximum likelihood approach, Chen & al. (2023) found full support for one *Pachycentria* species being nested among 24 *Medinilla* species. For a better understanding of the generic limits of *Medinilla*, and to verify the relationship and make necessary taxonomic changes, additional samples of *Pachycentria* are needed.

*Plethiandra* consists of eight species in Malesia, mostly confined to Borneo (Kadereit, 2005). It is easily recognized by its 6-merous flowers, polystaminate androecium, and inappendiculate anthers. Initially, *Plethiandra* was placed in the tribe Astronieae (Hooker, 1867), largely because of longitudinal anther dehiscence. Cogniaux (1891) maintained this association, but classified several new species that now belong to this group as *Medinillopsis* and *Medinilla robusta*. Stapf (1895) recognized the close affinity of *Plethiandra* to *Medinilla* and concluded the original anther description was misrepresentative. Taxonomists have continued to tentatively maintain the distinction (Kaderiet, 2005), but *Plethiandra* is still often accidentally included in *Medinilla*. For example, during the sorting of undetermined specimens of *Medinilla* from Borneo, *Plethiandra* is inevitably intermixed (Quakenbush, pers. obs.); and the "*Medinilla* sp. nov. Lin681" in Zhou & al. (2022) is *P. robusta* (Cogn.) Nayar (Fig. 7F). So far, molecular insights have been limited. Clausing & Renner (2001) did not resolve the relationship between six *Medinilla*, two *Plethiandra*, and 17 other Sonerileae. Maurin & al. (2021) found *P. robusta* sister to four *Medinilla*, with full support, based on genomic data. It remains to be seen whether this relationship to *Medinilla* will hold with greater sampling.

*Heteroblemma* was revised and established as a genus by Cámara-Leret & al. (2013). Previously, it was treated as a section of *Medinilla* (Blume, 1849). A total of 15 species are recognized in Vietnam and Malesia (POWO, 2024). *Heteroblemma* is generally characterized by a wood stele that is lobed in transverse section (Fig. 6B; also see Cámara-Leret & al. 2013), alternate leaves (via strong anisophylly and abortion) with prominent transverse venation, sessile and fascicled flowers along the stem, isomorphic stamens, woody berries, and papillate seeds. Molecular data show *Heteroblemma* and *Medinilla* in separate clades (Zhou & al., 2019; Zhou & al., 2022). In a plastome phylogeny (Zhou & al., 2022, Fig. S6), three *Heteroblemma* and four *Phyllagathis* Blume species formed a mixed clade sister to eight *Medinilla*. In a nuclear genomic phylogeny (Zhou & al., 2022), the same *Heteroblemma* accessions were monophyletic and part of a clade including nine *Phyllagathis* and seven *Driessenia* Korth. species. Notably, they were not sister to *Medinilla*. Despite this discordance, sampled *Heteroblemma* and *Medinilla* were clearly separate in both cases.

*Catanthera* was revised by Nayar (1982). It also has a long association with *Medinilla*; for example, Mansfeld (1926) transferred some members of *Hederella* Stapf (later synonymized as *Catanthera*) to *Medinilla*, and Bakhuizen van den Brink, Jr. (1943) thought it should be a section of *Medinilla*. *Catanthera* includes 19 species restricted to Malesia (POWO, 2024). It shares the atypical wood anatomy of *Heteroblemma* and is generally distinguished by opposite or alternate leaves with more obscure transverse venation; axillary or cauliflorous, umbellate or paniculate inflorescences; isomorphic or dimorphic stamens; soft, juicy berries; and smooth seeds. Clausing & Renner (2001) found weak support for a close relationship between *Catanthera* and *Heteroblemma* based on evidence from three chloroplast markers. Subsequent studies have not sampled these taxa to verify this relationship, but their close association has been accepted due to morphological similarity and shared geography (e.g., Cámara-Leret & al., 2013; Liu & al., 2022).

*Kendrickia* is enigmatic, monotypic, and most likely only found in Sri Lanka. Though it has been reported from the Anamala Hills (Clarke, 1879) and South India more generally (Triana, 1871; Bremer & Lundin, 1988), no specimens have been found to verify this. Like *Heteroblemma* and *Catanthera*, it has a lobed stele in transverse section and a climbing habit. It is distinguished by opposite, isophyllous leaves with obscure transverse venation; terminal or axillary inflorescences; isomorphic stamens; fleshy capsules that rupture at maturity; and smooth, prism-shaped seeds. Despite the unique morphological traits (e.g., fruit and seed type) and a geographic distribution outside of Malesia, Clausing & Renner (2001) found support (BS = 94) for a *Catanthera* and *Kendrickia* clade sister (BS = 50) to *Heteroblemma* based on sequences from a single chloroplast marker (*ndhF*). This close relationship has been accepted by subsequent authors (e.g., Cámara-Leret & al., 2013; Liu & al., 2022). However, this limited molecular evidence needs verification.

In light of the evidence, Liu & al. (2022) divided the fleshy-fruited Sonerileae into two alliances. The *Medinilla*-alliance includes *Medinilla*, *Pachycentria*, and *Plethiandra*. It is characterized by typical wood anatomy (i.e., terete stele) and soft, juicy berries. The monophyly of *Medinilla* is in question, and its major lineages are poorly understood. The *Heteroblemma*-alliance includes *Heteroblemma*, *Catanthera*, and *Kendrickia*, characterized by atypical wood anatomy (i.e., lobed stele) and either woody berries, soft, juicy berries, or fleshy capsules. While genera in this alliance are well-defined, their relationship to each other and the *Medinilla*-alliance requires further exploration. As such, the primary aims of this study are to explore the relationships among the fleshy-fruited Sonerilean genera, test the monophyly of *Medinilla*, identify major lineages within *Medinilla*, and update the taxonomy to reflect natural groups.

## MATERIALS AND METHODS

### Sampling, DNA extraction, library preparation, target capture, and sequencing

Silica dried plant material and (rarely) herbarium material were targeted from species belonging to the formal and informal groups discussed in the introduction and major clades identified by Zhou & al. (2019, 2022). Species from major biogeographic realms where *Medinilla* species naturally occur were also targeted, namely the Afrotropical, Indomalayan, Australasian, and Oceanian realms of Olson & al. (2001). Total genomic DNA was extracted using a modified CTAB method (Majure & al., 2019). Instead of isolating a pellet of DNA, samples were directly cleaned with a QIAquick PCR purification kit (QIAGEN, Hilden, Germany). Quality and yield of the DNA samples were checked using a Qubit 4 (Thermo Fisher Scientific, Waltham, MA, USA).

Library preparation, target sequence capture, and genome skimming were performed by Rapid Genomics LLC (Gainesville, FL, USA), using their high-throughput sequencing workflow and proprietary chemicals. A set of Melastomataceae-specific probes (Jantzen & al., 2020) targeting 384 loci were used to hybridize with the library inserts. Paired-end reads (2 x 150 base pairs) were generated on the Illumina NovaSeq 6000 system (San Diego, CA, USA).

In addition to the 126 samples sequenced in this study, we utilized data from various sources. Raw sequencing data from 66 accessions from Maurin & al. (2021) were obtained from the European Nucleotide Archive (ENA) using enaBrowserTools. RNA-seq data for *Medinilla magnifica* (Leebens-Mack & al., 2019) were accessed from the NCBI Sequence Read Archive (SRA). Furthermore, whole genome resequencing data from 83 accessions from Zhou & al. (2019, 2022) were incorporated. For a comprehensive list of samples and voucher information, refer to Supplementary Table S1.

### Nuclear data processing and analysis

Raw reads of target capture and RNA-seq data were processed to remove adaptors, trim low-quality bases, trim polyG and polyX tails, and filter out very short reads with fastp (-g -x -r -l 30 -5 --cut_front_window_size 1 -3 _tail_window_size 1 -- detect_adapter_for_pe) (Chen & al., 2018). HybPiper (Johnson & al., 2016) was used to recover targeted nuclear loci using the updated Melastomataceae probe set (Dagallier & Michelangeli, 2024) as reference file. Diamond (Buchfink & al., 2015), instead of BWA (Li & Durbin, 2009), was employed for read mapping to have better recovery.

To extract reads from the whole genome resequencing data from Zhou & al. (2019, 2022), reads from 12 *Medinilla* species obtained via target capture in the present study were mapped to a draft genome of *Tigridiopalma magnifica* using BWA-MEM (Li & Durbin, 2010). The resulting BAM files were merged into one using SAMtools (Li & al., 2009). The overall depth of each position was calculated, and a BED file containing regions with depth > 600 (> 50 per sample) was output using BEDtools (Quinlan & Hall, 2010). Whole genome resequencing data were then mapped to the reference genome. Finally, reads aligned to the regions listed in the BED file were extracted, and sequence assembly was carried out using HybPiper as described above.

The HybPiper command “paralog_retriever” was run to retrieve all the gene copies of the recovered loci for orthology inference. The sequences of each locus were aligned using MAFFT v7.453 (Katoh & Standley, 2013) with the method L-INS-I to provide accuracy. Reverse complement sequences were generated, if necessary, and were aligned with other sequences using the function “--adjustdirectionaccurately”. Phyutility v2.7.1 (Smith & Dunn, 2008) was subsequently used to trim the alignments. Sites missing 90% or more data were deleted. Sequences with more than 90% gaps or that were too short (< 50 base pairs) were also removed. IQ-Tree v2.1.3 (Minh & al., 2020) was used to infer single gene trees with 1,000 ultrafast bootstrap replicates. If a taxon was represented by monophyletic or paraphyletic tips, single gene trees were trimmed to keep only the tips with the most unambiguous characters, following Yang & Smith (2014) and Morales-Briones & al. (2021). TreeShrink v1.3.7 (Mai & Mirarab, 2018) was used with default settings to remove abnormally long branches from the single gene trees and the corresponding sequences from the alignments.

DISCO-ASTRAL (Willson & al., 2021) was used to infer orthologs. DISCO takes gene trees as input, roots and labels the internal nodes as either duplication or speciation events using the method implemented in ASTRAL-Pro (Zhang & al., 2020), and decomposes the gene trees with its decomposition algorithm. DISCO was run with default settings except the option -m 20 was used for filtering trees with less than 20 tips. Finally, ASTRAL v5.7.8 (Zhang & al., 2018) was used with the decomposed gene trees as input to infer the species tree (nuclear tree). Branch support was quantified by Local Posterior Probability (LPP; Sayyari & al., 2016).

### Plastid loci assembly and phylogenomic analysis

Plastid loci were recovered from off-target reads of the target capture data using HybPiper. Annotated plastomes of *Medinilla magnifica* (MT043350) and the aforementioned 83 accessions representing all major clades within Sonerileae, coupled with 18 plastomes representing most Melastomataceae tribes treated by Penneys & al. (2022), were used to extract protein-coding genes for target file construction using a script developed by Zhang & al. (2020; https://github.com/Kinggerm/GetOrganelle/blob/master/Utilities/get_annotated_regions_fr om_gb.py). BWA was used for read mapping. Samples with less than seven genes recovered were excluded from further analysis. The HybPiper command “retrieve_sequences” was used to retrieve the recovered plastid sequences, which were combined with sequences from the annotated Sonerileae and Dissochaeteae plastomes. Sequences of each plastid gene were aligned in the same way as nuclear data. AMAS (Borowiec, 2014) was used to concatenate them into a supermatrix. A Maximum Likelihood (ML) phylogeny was estimated in IQ-Tree. The IQ-Tree option “-m MFP + MERGE” was used to select the best-fitting partition scheme and models before tree reconstruction. Coalescent analysis was not conducted for plastid data, as the whole plastome has been suggested to be treated as a coalescent gene (Doyle, 2021).

## RESULTS

### Phylogenetic analyses

The phylogeny of the fleshy-fruited Sonerileae was reconstructed in two different ways based on two different sets of data (Figs. 1–5). The final DISCO-ASTRAL tree based on nuclear sequence data (NAT; short “nuclear tree”) includes 272 samples (79 samples from Zhou & al. (2019, 2022), 1 sample from Leebens-Mack & al. (2019), 66 samples from Maurin & al. (2021), and 126 samples sequenced in the present study) and is based on 599 single gene trees generated by DISCO. For a summary of targeted genes recovered for each sample, refer to Supplementary Table S2. The partitioned ML concatenation tree based on 81 plastid protein-coding genes (PCT; short “plastid tree”) includes 224 samples (84 whole plastomes, 45/66 samples from Maurin & al. (2021); and 95/126 samples sequenced in the present study). Each sample is represented by at least seven loci (see Supplementary Table S2). The total combined sequence length of the plastid supermatrix is 69943 base pairs, 33.96% of which is gaps/missing data.

**Figure 1.**
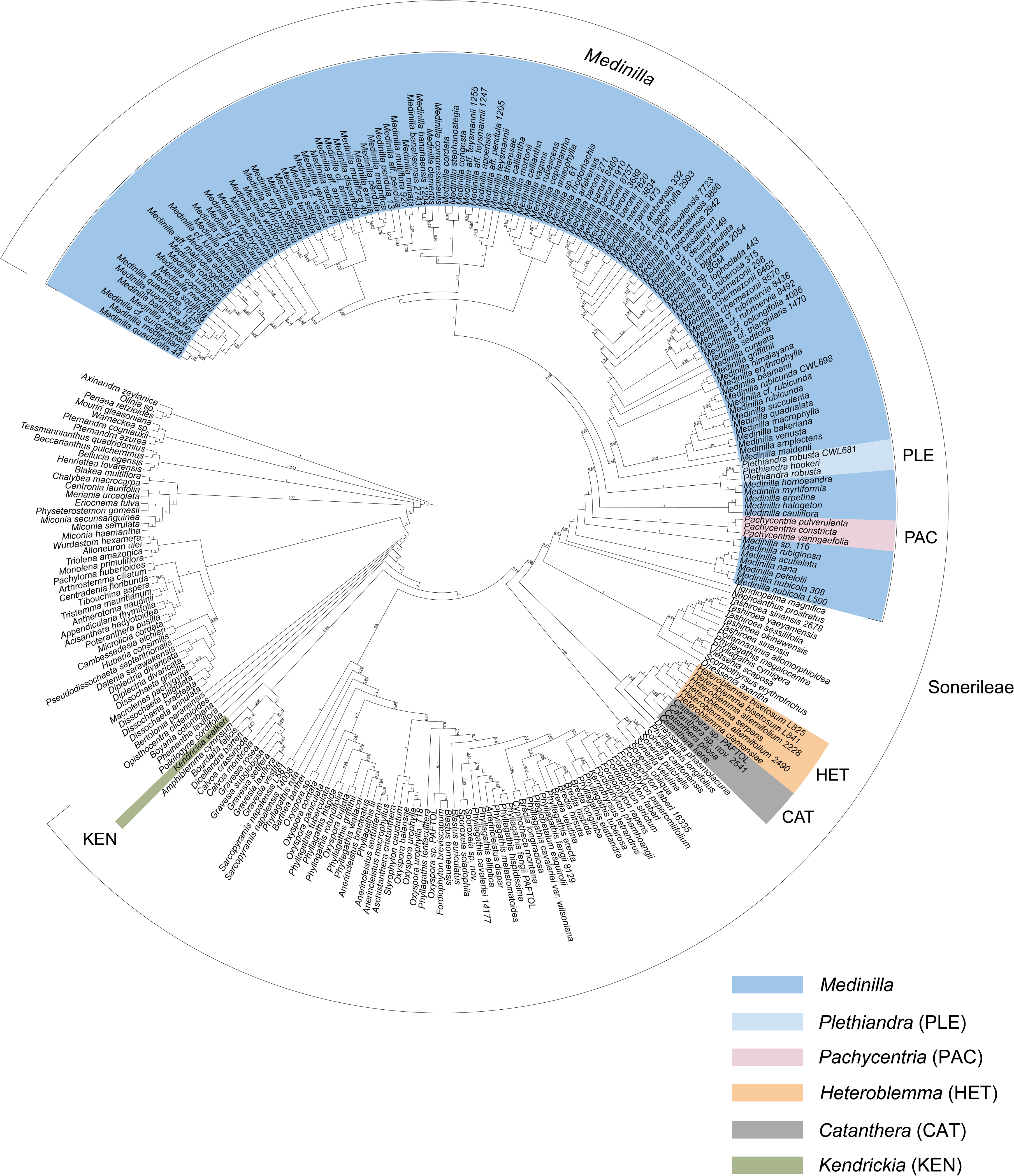
DISCO-ASTRAL tree based on 272 samples and 385 nuclear genes showing the relationships of the fleshy-fruited Sonerileae genera (colored ranges) in the context of Sonerileae (bracketed range). Local Posterior Probabilities (LLP) are shown above the branches.

Fleshy-fruited Sonerileae are resolved in three separate clades (Figs. 1 & 2). In the nuclear tree, *Kendrickia* appears as an early diverging lineage in Sonerileae, sister to the Afrotropical clade (NAT: LPP = 0.69). Notably, *Kendrickia* does not fall within the Asian Superclade to which the other fleshy-fruited genera belong (NAT: LPP = 1). Not enough sequences were recovered to include *Kendrickia* in the analysis of the plastid sequence data. In the nuclear tree (Fig. 1), four *Catanthera* and six *Heteroblemma* have full support as monophyletic sister clades. They belong to a clade that includes *Phyllagathis longifolius* and *Driessenia phasmolacuna* (NAT: LPP = 1). *Tigridiopalma* C.Chen is sister to *Medinilla* (NAT: LPP = 0.81). In the plastid tree (Fig. 2), *Heteroblemma* is not monophyletic. Four samples are sister to three *Catanthera*, while two samples are sister to *P. longifolius*. This *Catanthera*-*Heteroblemma*-*Phyllagathis* clade is sister to the *Medinilla* clade. *Driessenia phasmolacuna* is resolved in the sister group to these two clades, along with other *Driessenia* and *Phyllagathis* species. All of these relationships have strong support (PCT: BS = 98–100).

**Figure 2.**
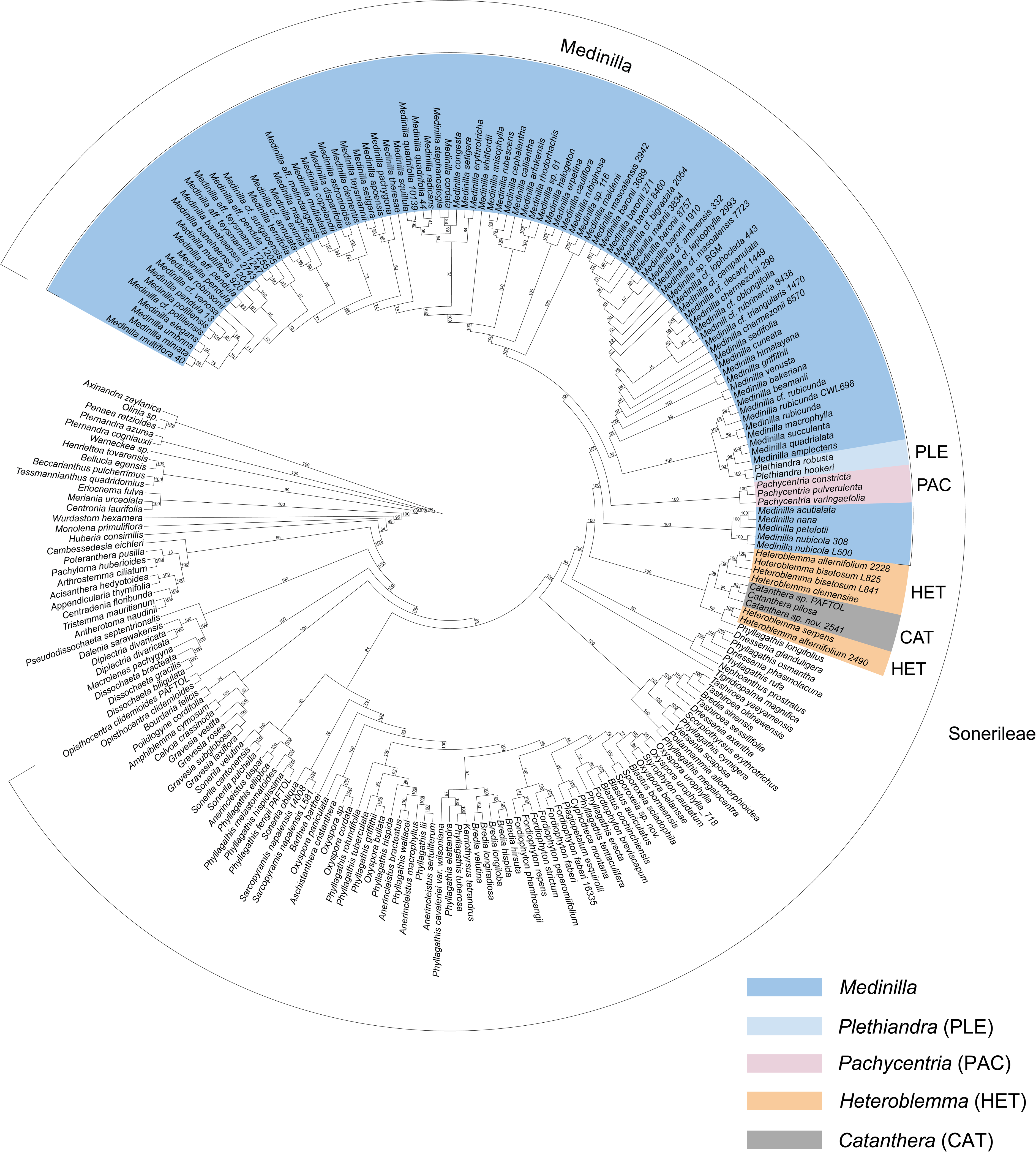
Maximum Likelihood plastid tree based on 81 genes and 224 samples showing the relationship of the fleshy-fruited Sonerileae genera (colored ranges) in the context of Sonerileae (bracketed range). Bootstrap (BS) support values are shown above the branches.

Both phylogenies resolve *Medinilla* as paraphyletic with respect to *Pachycentria* and *Plethiandra* (NAT: LPP = 1, PCT: BS = 100; Figs. 1–3). It includes all three sampled *Pachycentria* and all three sampled *Plethiandra* (Figs. 1–3). There are seven “Early Diverging Clades” (Fig. 3). The *M. nubicola*-alliance (NAT: LPP = 1, PCT: BS = 100; Fig. 3) is sister to all other *Medinilla* species sampled (NAT: LPP = 1, PCT: BS = 100). In the nuclear tree (Fig. 3, left), the *M. rubiginosa*-alliance diverges next (NAT: LPP = 1), followed by *Pachycentria* (NAT: LPP = 1). Then the *M. erpetina*-alliance diverges (NAT: LPP = 1), but the placement has low support (NAT: LPP = 0.61). The *M. myrtiformis*-alliance (NAT: LPP = 1) and *Plethiandra* (NAT: LPP = 1) form the next sister group (NAT: LPP = 1), but with low support (NAT: LPP = 0.54). Similarly, the placement of the *M. maidenii*-alliance has low support (NAT: LPP = 0.65). It is resolved as sister to the Western Superclade (Fig. 4, left; NAT: LPP = 0.98). The Western Superclade consists of the *M. rubicunda*-alliance (NAT: LPP = 0.95), *M. erythrophylla*-alliance (NAT: LPP = 1), *M. cuneata*-alliance, *M. sedifolia*-alliance, and *M. viscoides*-alliance (NAT: LPP = 1). The Eastern Superclade (Fig. 5, left; NAT: LPP = 1) consists of three clades: the *M. arfakensis*-alliance (NAT: LPP = 1) and *M. anisophylla*-alliance (NAT: LPP = 1) are sister groups (NAT: LPP = 1) and together are sister to the *M. medinilliana*-alliance (NAT: LPP = 1).

**Figure 3.**
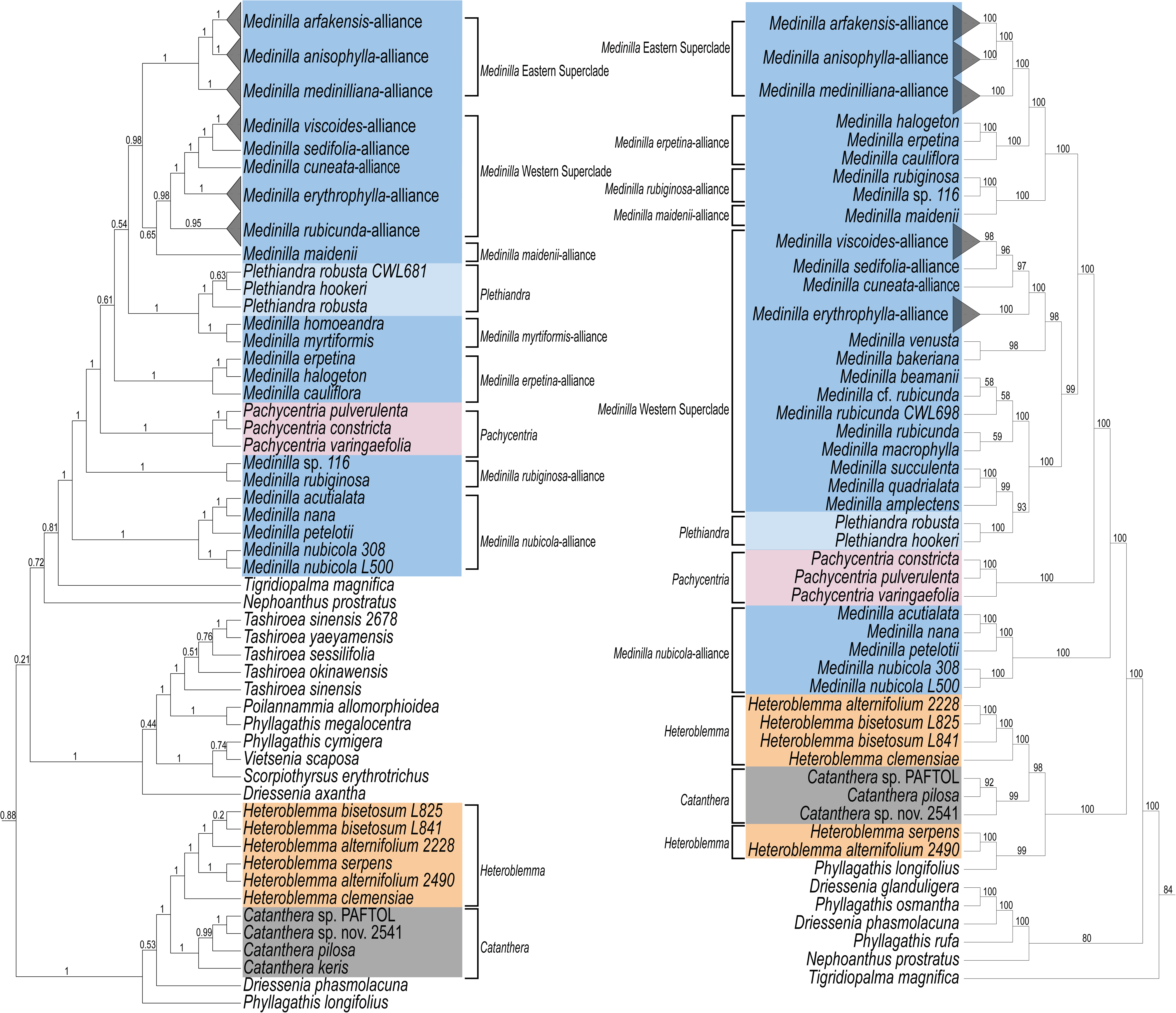
DISCO-ASTRAL nuclear tree (left) based on 385 genes and 272 samples and Maximum Likelihood plastid tree (right) based on 81 genes and 224 samples showing the Early Diverging Clades of *Medinilla* and sister taxa. Local Posterior Probability (nuclear tree) and Bootstrap support values (plastid tree) are shown below the branches

**Figure 4.**
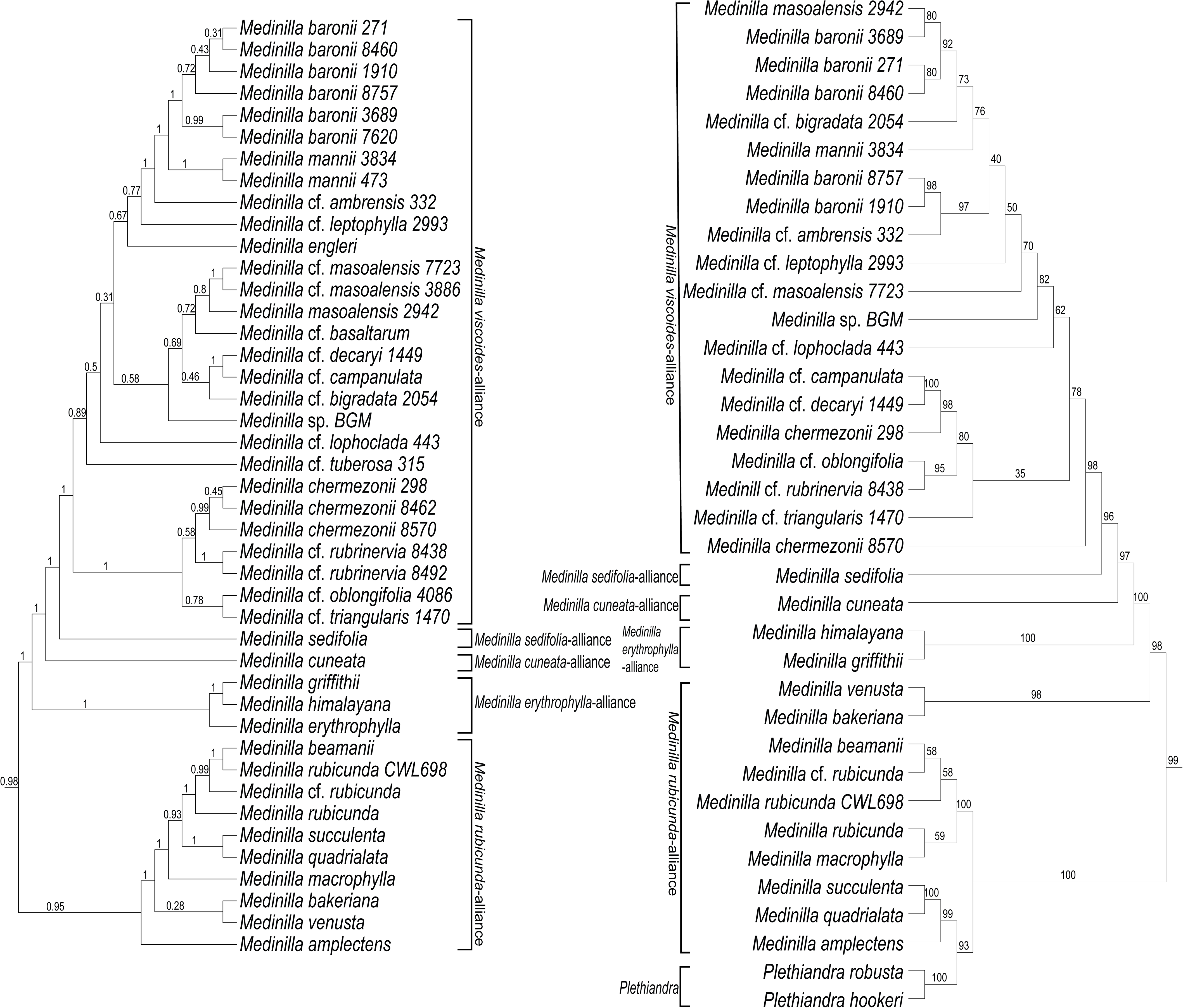
Alliances within the Western Superclade of *Medinilla* are shown on an DISCO-ASTRAL nuclear tree (left) based on 385 genes and 272 samples and a Maximum Likelihood plastid tree (right) based on 81 genes and 224 samples. Local Posterior Probability (nuclear tree) and Bootstrap support values (plastid tree) are shown below the branches

Most of the major *Medinilla* clades recovered in the nuclear tree (Figs. 3–5, left) were also strongly supported in the plastid tree (Figs. 3–5, right), but their relationships to each other often differ, especially in the case of the Early Diverging Clades (Fig. 3). *Pachycentria* diverges after the *M. nubicola*-alliance (PCT: BS = 1). The *M. maidenii*-alliance is sister (PCT: BS = 100) to the *M. rubiginosa*-alliance (PCT: BS = 100). These are sister (PCT: BS = 100) to the *M. erpetina*-alliance (PCT: BS = 100) and the Eastern Superclade (PCT: BS = 100). *Plethiandra* is resolved among species from the *M. rubicunda*-alliance (PCT: BS = 100), which is resolved in two separate clades (PCT: BS = 99). There are many inconsistencies between the species relationships of the *M. viscoides*-alliance. Similarly, there are many inconsistencies within the *M. medinilliana*-alliance (Fig. 5).

**Figure 5.**
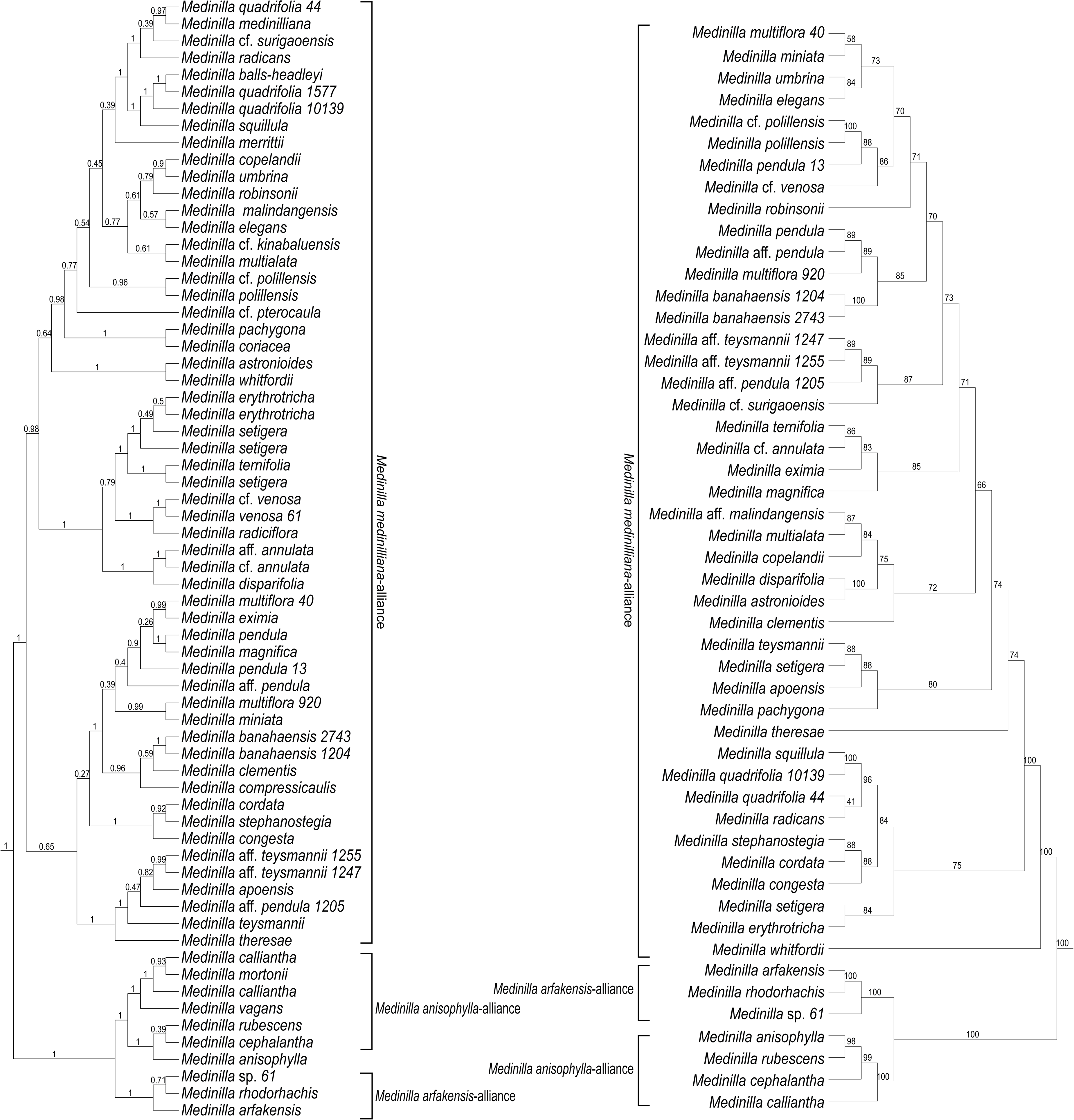
DISCO-ASTRAL nuclear tree (left) based on 385 genes and 272 samples and Maximum Likelihood plastid tree (right) based on 81 genes and 224 samples showing the Eastern Superclade of *Medinilla.* Local Posterior Probability (nuclear tree) and Bootstrap support values (plastid tree) are shown below the branches

## DISCUSSION

Our phylogenetic study representing 227 accessions of Sonerileae, including 141 accessions of *Medinilla* and allies, revealed that the fleshy-fruited Sonerileae belong to at least three different clades within the tribe: 1. *Kendrickia*, 2. *Heteroblemma/Catanthera*-alliance, and 3. *Medinilla* (including *Pachycentria* and *Plethiandra*). *Kendrickia* is resolved in an isolated phylogenetic position and is not part of the *Heteroblemma/Catanthera*-alliance. *Pachycentria* and *Plethiandra* are nested within *Medinilla*. Each clade is discussed below. For *Medinilla*, fifteen major lineages were identified and are further discussed below. These lineages are ordered based on their appearance in the nuclear tree (Fig. 3, left), indicating the estimated divergence from the *Medinilla* type species (*M. medinilliana*). The observation of widespread discordance between the nuclear and plastid trees is discussed in the respective sections.

### Kendrickia

*Kendrickia* (Fig. 6A) is not closely related to any of the other fleshy-fruited, climbing Sonerileae with lobed steles. Thus, fleshy fruit and atypical xylem evolved convergently in Sonerileae, the latter probably in relation to the root climbing habit (Cámara-Leret & al., 2013). Rather, *Kendrickia* is sister to the Afrotropical clade of Sonerileae in the nuclear tree (Fig. 1; see Liu & al., 2022; Veranso-Libalah & al., 2022), albeit with low support (NAT: LPP = 0.69). Plastid sequences from *Kendrickia* were not included in the analysis, so this relationship could not be re-tested. However, there are some noteworthy morphological similarities to the Afrotropical Superclade. For instance, some *Gravesia*, *Dicellandra*, and *Calvoa* are reported to be root-climbers (Veranso-Libalah & al., 2022). Additionally, the first two have pyramidal and wedge-shaped seeds, respectively, somewhat resembling the prism-shaped seeds of *Kendrickia*. Furthermore, fruit dehiscence in *Dicellandra* is somewhat akin to *Kendrickia*, via the rupturing of the capsule wall. The fleshy capsules of *Kendrickia* rupture when ripe and have only a superficial resemblance with the true berries of *Medinilla*, *Catanthera*, and *Heteroblemma*. These results raise new questions about *Kendrickia*’s phylogenetic relationships and geographic origins.

**Figure 6.**
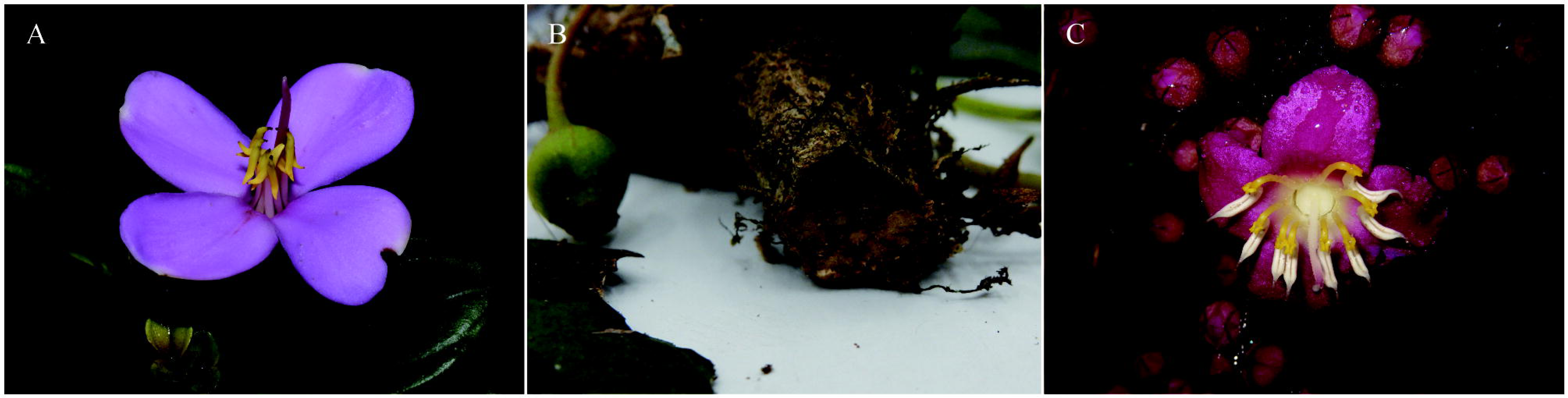
Fleshy-fruited Sonerileae with atypical xylem configuration. (A) *Kendrickia* flower and leaves (Photo: Bathiya Gopallawa, KED 01, Sri Lanka); (B) *Catanthera sp.* fruit and stem cross-section with lobed xylem cross-section (Photo: Darin Penneys 2523, Borneo); (C) *Heteroblemma clemensiae* flowers and buds (Photo: Maxim Nuraliev 1345, Vietnam).

### Catanthera *and* Heteroblemma

The close relationship between *Catanthera* (Fig. 6B) and *Heteroblemma* (Fig. 6C) is robustly supported, forming a clade with full support in the nuclear tree (Fig. 3, left). However, the plastid tree (Fig. 3, right) presents a more complex picture. Our study represented the first inclusion of *Cantanthera* in phylogenomic study and included twice as many *Heteroblemma* samples as Zhou & al. (2022), and the same relationship with *Phyllagathis* was resolved with full support. Specifically, *Heteroblemma* from Vietnam are sister to *Catanthera*, while *Heteroblemma* from Malesia are part of a clade that includes capsular-fruited *Phyllagathis* species. Despite this, given *Heteroblemma’s* morphological cohesion and strong support from the nuclear tree, it is still reasonable to treat it as a distinct genus. The topology of both trees supports the continued recognition of *Catanthera*.

The relationship of *Catanthera* and *Heteroblemma* to *Medinilla* is conflicting, similar to the results of previous studies (Zhou & al., 2019; Zhou & al., 2022). In our plastid tree (Figs. 2 & 3), *Catanthera* and *Heteroblemma* are part of a sister clade to *Medinilla* that includes a *Phyllagathis* species with thyrsoid inflorescences. Despite different taxon sampling, Zhou & al. (2022) also identified a similar clade, comprising *Phyllagathis* with thyrsoid inflorescences and *Heteroblemma*, as being closely related to *Medinilla*. In our nuclear tree (Figs. 1 & 3), *Tigridiopalma* is sister to *Medinilla*, together forming a clade sister to *Nephoanthus* C.W.Lin & T.C.Hsu. In comparison, Zhou & al. (2022) discovered a clade that combines *Tigridiopalma* and *Nephoanthus* as a sister group to *Medinilla*. To account for the observed widespread phylogenetic discordance in Sonerileae, Zhou & al. (2022) identified several contributing factors, including random noise from uninformative genes, incomplete lineage sorting (ILS), and hybridization or introgression. It is worth noting that branch lengths are very short in this area (Supplementary Fig. S3 & S4). and the discord may not be real. Whatever the case, these relationships are still not well understood and warrant further exploration. However, as with *Kendrickia*, it is probable that fleshy fruit evolved independently in this clade as well.

### Medinilla Overview

The limits of *Medinilla* were rigorously tested, incorporating all major taxonomic groups and alliances either directly or indirectly via similar/associated taxa. Species were sampled from across the geographic range of *Medinilla*, spanning from West Africa to the Solomon Islands, and from China to Australia, encompassing the four greatest centers of species diversity: Madagascar, Borneo, the Philippines, and New Guinea. Notably, both *Pachycentria* and *Plethiandra* are nested among *Medinilla* species (Figs. 1–3). With the inclusion of these two genera, *Medinilla* can be distinguished from all other taxa in Sonerileae by the combination of typical xylem (vs. lobed in cross-section) and berry fruit. While these similar traits are also found in the Dissochaeteae, they can be distinguished by alternate inter-vessel pits (as opposed to being scalariform in *Medinilla*; see van Vliet, 1981), interpetiolar ridges and generally more chartaceous leaves with basal acrodromous venation. Phylogenetically, *Medinilla* can be identified as the most exclusive clade containing *M. medinilliana* and *M. nubicola*. Major clades are divided into three groups: the Early Diverging Clades, the Western Superclade, and the Eastern Superclade.

The Early Diverging Clades (Fig. 3) include some members that were previously considered as separate genera, such as *Pachycentria*, *Erpetina*, and *Plethiandra*. Each of these genera is characterized by atypical anthers, usually lacking ventral and sometimes dorsal appendages. They also encompass species from *M.* sect. *Heteromedinilla* and the *M. suberosa*-alliance (Clausing, 1999). The species within these clades range from the lower Himalayas to Vanuatu.

The Western Superclade primarily consists of species found west of the Wallace Line (for clarification, see Ali & Heaney, 2021). Major clades within this superclade diverge successively as the sampling progresses further west. The *Medinilla rubicunda*-alliance, primarily from Sundaland, includes species placed in *M.* sect. *Sarcoplacuntia*, *M.* sect. *Apateon*, *M.* sect. *Heteromedinilla*, and various informal alliances treated by Regalado (1990, 1995) and Clausing (1999). The *M. erythrophylla*-alliance, from mainland Asia (and Hainan), includes a species placed in the *M. suberosa*-alliance (Clausing, 1999). *Medinilla cuneata* is found in Sri Lanka, and shares similarities with other species in the region. Meanwhile, the *M. sedifolia* and *M. viscoides* alliances are from the Afrotropical realm, primarily Madagascar. *Triplectrum* is associated with the *M. sedifolia*-alliance, and *Diplogenea*, *M.* sect. *Septatae*, *M.* sect. *Adhaerentes*, and Clausing’s (1999) Group 2 are associated with the *M. viscoides*-alliance.

The Eastern Superclade primarily comprises samples collected east of Huxley’s modification of the Wallace Line (i.e., the Huxley Line; see Ali & Heaney, 2021). This superclade is split into two major clades. One consists of samples from New Guinea, the Bismarck Archipelago, and the Solomon Islands (the *Medinilla arfakensis* and *M. anisophylla* alliances). The other clade (the *M. medinilliana*-alliance) is mainly composed of Philippine species. Nested within these clades are samples from other regions in the Indomalayan, Australasian, and Oceanian realms. Notably, the *Medinilla* type species (*M. medinilliana*) belongs to this clade, along with species once considered *Dactyliota*, *Hypenanthe*, and *Carionia*. *Cephalomedinilla* is also associated with this clade. Additionally, several *M.* sect. *Sarcoplacuntia* and *M.* sect. *Heteromedinilla* species, all species sampled from *M.* sect. *Medinilla*, and many species sampled from the informal alliances treated by Regalado (1990, 1995) and Clausing (1999; Group 1) were resolved within this superclade.

### Medinilla nubicola-alliance

The relationship of *Medinilla nubicola* (= *M. fengii*) and *M. petelotii* (Fig. 7A) was initially established by Zhou & al. (2019, 2022), along with their sister position to a few other *Medinilla* species. In this study, two additional associates were identified, and their sister relationship to the rest of *Medinilla* is fully supported by both nuclear and plastid trees. Notably, some species in this clade were originally associated with *Pachycentria*. For example, *P. formosana* Hayata and *P. fengii* S.Y.Hu are synonyms of *M. nubicola*. *Medinilla nana* was compared to *P. fengii* in the protologue (Hu, 1952), and *M. arunachalica* G.D.Pal (not sampled but very similar to *M. nana*) was compared to *M. maingayi* C.B.Clarke (Pal, 1995), a subspecies of *P. glauca*. Clausing (2000) excluded *M. nubicola* from *Pachycentria* based on seed morphology, a distinction supported here. In this alliance, anthers are falcate, with long or nearly absent (*M. nubicola*), needle-like, ventral appendages and a tapered dorsal spur (Fig. 7A). These species are found in mainland Asia and Taiwan.

**Figure 7.**
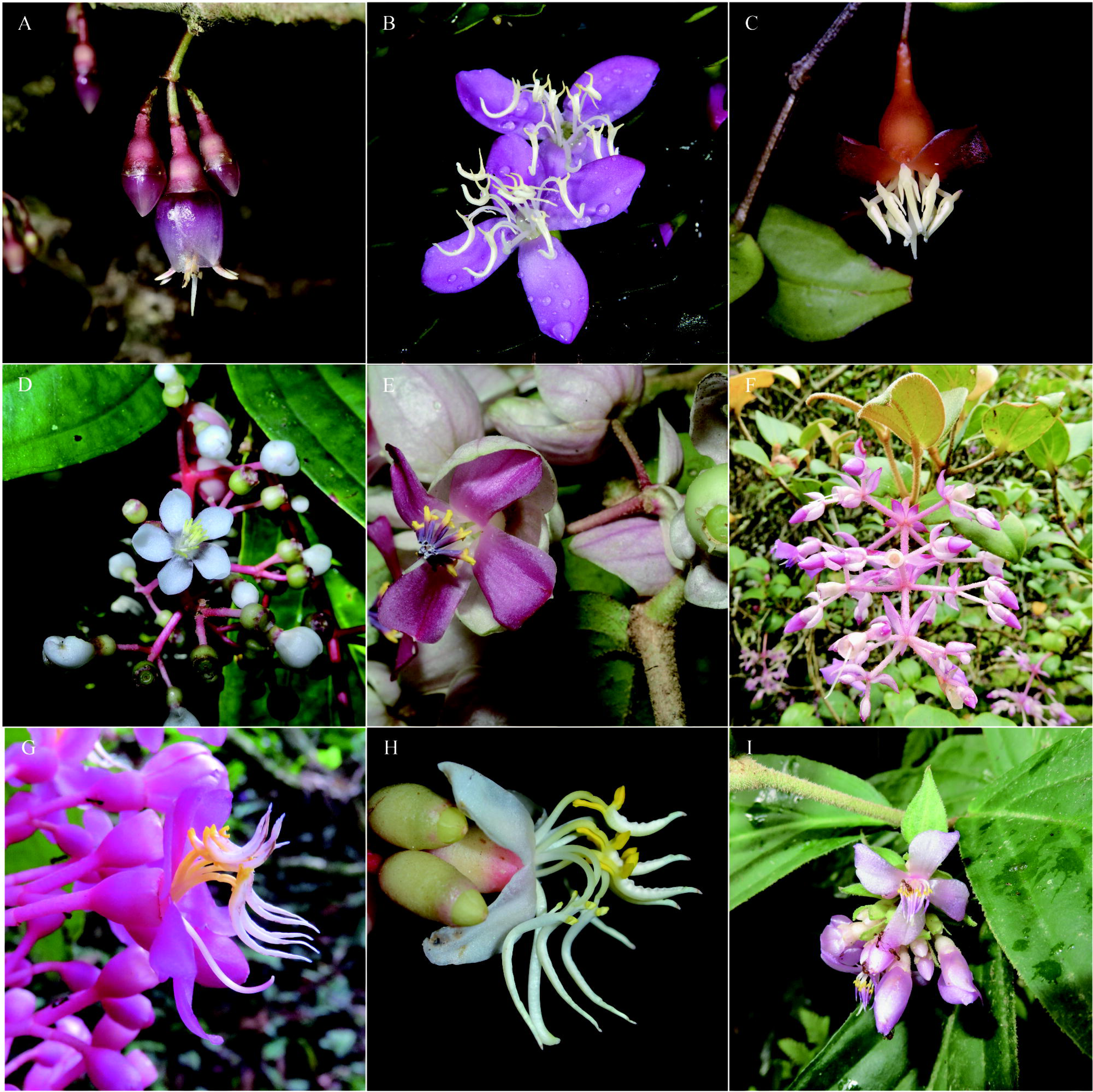
Representatives from major lineages identified within *Medinilla*: (A) *M. petelotii* in the *M. nubicola*-alliance (Photo: Maxim Nuraliev, Vietnam); (B) *M. rubiginosa* in the *M. rubiginosa*-alliance (Photo: Barry Conn 5361, New Guinea); (C) *M. pulverulenta* in *Pachycentria* (Photo: Selby Gardens 2016-0223A, in cultivation, origin unknown); (D) *M. erpetina* in the *M.* e*rpetina*-alliance (Photo: Patrick Blanc, Solomon Islands); (E) *M. myrtiformis* in *M. myrtiformis*-alliance (Photo: PLSPH 807, Philippines); (F) *M. robust*a in *Plethiandra* (Photo: Che-Wei Lin 681, Borneo); (G) *M. maidenii* in the *M. maidenii*-alliance (Photo: Selby Gardens, MSBG2002-0198B, in cultivation, originally from New Guinea); (H) *M. rubicunda* in the *M. rubicunda*-alliance (Photo: Maverick Tamayo, AVAMR 16, Philippines); and (I) *M. griffithii* in the *M. erythrophylla-*alliance (Photo: Kate Armstrong 2903, Myanmar).

### Medinilla rubiginosa-alliance

Two species from New Guinea were resolved in a fully supported clade by both datasets (Fig. 3). However, placement varies between the datasets. According to the nuclear tree (Fig. 3, left), the *Medinilla rubiginosa*-alliance diverged after the *M. nubicola-* alliance. Conversely, in the plastid tree (Fig. 3, right), the *M. rubiginosa*-alliance is associated with *M. maidenii* (Fig. 7G; see *M. maidenii*-alliance) and is sister to the *M. erpetina*-alliance and the Eastern Superclade. All of these clades overlap in distribution and are found east of the Wallace Line.

Shared traits of the *Medinilla rubiginosa*-alliance include hairiness, leaves with nerves arising near the base, terminal inflorescences, conspicuous bracts, and somewhat verrucose berries (Fig. 7B). In the case of *M. rubiginosa*, anthers have three, relatively equal appendages pointing basally. Anther details of *Medinilla* sp. 116 are unknown.

Clausing (2000) excluded *Medinilla rubiginosa* (Fig. 7B; = *Pogonanthera hexamera* Baker f.) from *Pachycentria*, a decision fully supported in this study. It has not been associated with any other taxonomic groups. However, *M. pulleana* is a similar species from New Guinea that was transferred to *Hypenanthe* by Bakhuizen van den Brink, Jr. (1943) because of its hairiness and conspicuous bracts. Clausing (1999) also placed *M. pulleana* with *Hypenanthe* species, but in its own subgroup, noting its closer resemblance to other New Guinea species than those of *Hypenanthe*. Subsequently, Bodegom & Veldkamp (2001) characterized a group of pseudostipular species from New Guinea and the Bismarck Archipelago. While the presence of pseudostipules in *M. pulleana* is ambiguous, it closely resembles the group in every other respect. Similarly, both species in the *M. rubiginosa*-alliance lack definite pseudostipules, but they share other similarities with the pseudostipular group. Bodegom & Veldkamp (2001) postulated that the pseudostipular species belong to a larger group including non-pseudostipular species, and a close relationship between the *M. rubiginosa*-alliance and the pseudostipular species is expected but requires further verification.

### Pachycentria

Nuclear and plastid trees (Fig. 3) consistently support the monophyly of *Pachycentria* as circumscribed by Clausing (2000), and its placement within *Medinilla*. However, placement of *Pachycentria* within *Medinilla* varies between datasets. In the nuclear tree (Fig. 3, left), *Pachycentria* diverges after the *M. rubiginosa*-alliance, while in the plastid tree (Fig. 3, right), it diverges after the *M. nubicola-*alliance. Despite limited sampling (three out of eight species), the inclusion of the type species (*P. constricta*) and the two most morphologically divergent taxa (*P. varingaefolia* and *P. pulverulenta* [Fig. 7C]) ensured robust testing. Notably, *P. varingaefolia* with its remarkably large flowers and dimorphic stamens with ventral appendages, was resolved as sister to the other two species. On the other hand, *P. pulverulenta* (≡ *Pogonanthera pulverulenta*) has auriculate leaf bases and a tuft of hairs instead of a dorsal appendage. Despite these morphological differences, they share the diagnostic traits of this clade, i.e., small ovary in a strongly constricted, urceolate hypanthium, and seeds with comb-shaped testa cells.

### Medinilla erpetina-alliance

*Erpetina* (Fig. 7D) was established by Naudin in 1851 and later transferred to *Medinilla* by Triana in 1871. This move receives robust support from molecular data. *Medinilla erpetina* forms part of a three-species clade, with full support from both molecular datasets (Fig. 3). In the nuclear tree (Fig. 3, left), this clade diverged after *Pachycentria*, while in the plastid tree (Fig. 3, right), it is sister to the Eastern Superclade. Indeed, all three species within this clade are found east of the Wallace Line, in the Bismarck Archipelago, Solomon Islands, and Vanuatu.

Species within this clade are characterized as epiphytic shrubs or climbers with few-flowered, axillary inflorescences. They possess anthers with a prominent dorsal appendage and no ventral appendages. Clausing (1999) placed *Medinilla cauliflora* and *M. halogeton* within the *M. suberosa*-alliance, a combination and expansion of Regalado’s *M. succulenta* (1990) and *M. palawanensis* alliances (Group 9; 1995). However, other members of the *M. suberosa*-alliance tested in our study (*M. amplectens*, *M. erythrophylla*, *M. multialata*, and *M. succulenta*) were resolved in the *M. rubicunda* and *M. erythrophylla* alliances. Anther details for *M. palawanensis* are insufficiently known, but a more recently described species and presumed close relative, *M. ultramaficola*, has anthers somewhat consistent with those of the *M. erpetina*-alliance, having a long dorsal appendage and essentially lacking ventral appendages (Quakenbush & al., 2020). More sampling from Palawan is needed to better understand this group.

### Medinilla myrtiformis-alliance

The two species resolved in this clade were initially classified in genera other than *Medinilla*. *Medinilla myrtiformis* (Fig. 7E) was first described as *Aplectrum* Blume (= *Diplectria* [Blume] Rchb., tribe Dissochaeteae) and later transferred to *Medinilla* by Triana (1871). It is also synonymous with *Kibessia celebica* Miq. Similarly, *M. homoeandra* was initially classified as *Anplectrum* A.Gray (= *Diplectria*), and later transferred to *Medinilla* by Nayar (1966). These taxonomic reclassifications find full support from molecular evidence in both the nuclear and plastid trees (Fig. 3).

Both species are part of the previously identified *Medinilla myrtiformis*-alliance, a group characterized by shared traits that have been recognized for a long time (e.g., notes on *M. cardiophylla* in Merrill, 1910). Veldkamp (1978, 1988) undertook a revision of the group, with further contributions from Regalado (1990, 1995) and Clausing (1999). The defining features of this alliance are anthers that lack ventral appendages and have a short, triangular, dorsal plectrum. Additional shared traits include very narrow, divaricate branches; few-flowered, axillary cymes; and ovate-lanceolate petals. While *M. muricata* was initially placed within this group by Regalado (1990), it differs from typical members by a few key traits, such as petal shape (rounded vs. pointed) and anther appendages (ventral lobes present vs. absent). Recognizing these distinctions, Clausing (1999) placed *M. muricata* in a separate alliance. Flowers remain unknown for several other species assigned in this group, including *M. salicina*, *M. benguetensis*, and *M. gracilis*. Therefore, further investigations and testing are necessary to inform our understanding of these species.

*Medinilla homoeandra* is native to Borneo, while *M. myrtiformis* is found in Wallacea which includes the Philippines (see Ali & Heaney 2021). The alliance, including some unsampled species like *M. ericoidea* in New Guinea, demonstrates a widespread distribution in Malesia, spanning the Wallace Line. This distribution pattern aligns with other alliances, such as *Pachycentria*, the *M. rubicunda*-alliance, and the *M. medinilliana*-alliance. In the nuclear tree (Fig. 3, left), the *M. myrtiformis*-alliance was resolved as sister to *Plethiandra*. Unfortunately, too few sequences were recovered to include this alliance in the plastid tree and confirm this relationship.

### Plethiandra

*Plethiandra* is distinguished by its six-petaled flowers, more than double that number of stamens, and anthers without appendages. Molecular data fully supports the inclusion of *Plethiandra* in *Medinilla* (Figs. 1–4). The monophyly of *Plethiandra* is also well-supported. Although only the type, *P. hookeri*, and two accessions of *P. robusta* were sampled, representing two out of eight species, they were resolved together. *Plethiandra* stands out as one of the most easily distinguished groups, and its monophyly has never been in question (Kadereit, 2005). However, placement of *Plethiandra* within *Medinilla* varies depending on the dataset. In the nuclear tree (Fig. 3, left) it appears as a sister to the *M. myrtiformis*-alliance, positioned among the Early Diverging Clades of *Medinilla*. The reduced anther appendages observed in the *M. myrtiformis*-alliance, along with the absence of appendages in *Plethiandra*, suggest a potential affinity. Yet no samples of the *M. myrtiformis*-alliance were included in the plastid tree to verify this relationship. According to the plastid tree (Fig. 3, right), *Plethiandra* is nested among other Bornean taxa of the *M. rubicunda*-alliance.

### Medinilla maidenii-alliance

The classification of *Medinilla maidenii* (Fig. 7G) from New Guinea has long been uncertain. Mueller (1886) expressed no objection to placing it in *Pachycentria* due to atypical anthers for *Medinilla*. Anthers lack ventral appendages and possess a blunt dorsal projection. He also drew parallels with *Pternandra*, because the thecae are separated giving the appearance of dehiscence via a slit. Molecular evidence clearly places *M. maidenii* within *Medinilla*, but its internal placement remains ambiguous and in need of further exploration. In the nuclear tree (Fig. 3, left), it was resolved with low support as sister to the Western Superclade (NAT: LPP = 0.65), while in the plastid tree (Fig. 3, right), it was placed with full support as sister to the *M. rubiginosa*-alliance. The plant habit aligns more with the Western Superclade, whereas the geographic distribution aligns more with the *M. rubiginosa*-alliance. Notably, the anther morphology does not resemble either group. Kartonegoro (2022) synonymized three taxa from New Guinea with *M. maidenii*, and several additional taxa from New Guinea share a similar growth form and inflorescences (e.g., *M. nabirensis* and *M. papulosa*). Targeting these would help better understand the group.

### Medinilla rubicunda-alliance

Ten samples were resolved in a well-supported clade in the nuclear phylogeny (NAT: LPP = 0.95; Fig 4, left), originating from Sundaland (nine) and New Guinea (one), representing a sister group to species found further west. In the plastid tree, eight of these samples were resolved together (PCT: BS = 100), but *Plethiandra* is also included. The remaining two species, *Medinilla venusta* King and *M. bakeriana* Mansf., are resolved together in the sister group.

Members of the *Medinilla rubicunda*-alliance are epiphytic shrubs with often warty stems and glabrous nodes. Inflorescences lack persistent or conspicuous bracts and bracteoles, and they exhibit variable architectures. In paniculate inflorescences, branches are not arranged in a regular whorled pattern. The hypanthium is glabrous; anthers are equal, isomorphic, and possess two ventral lobes and generally a small dorsal spur. This alliance shares similarities with the *M. erythrophylla*-alliance, which is part of its sister group. Phylogenetic estimates suggests that previous classifications of these species (Blume 1831; Blume 1949; Bakhuizen, 1943; Regalado 1990; Regalado, 1995; Cluasing, 1999) are either polyphyletic or insufficient to capture the diversity within the *M. rubicunda*-alliance. Targeted sampling from Sundaland would likely help fill in gaps related to this group.

In the nuclear tree (Fig. 4, left), *Medinilla beamanii* was resolved within a clade comprising *M. rubicunda* samples. These species share strong morphological similarities with *M. beamanii*, differing primarily in having a longer peduncle and more umbellate inflorescences. *Medinilla rubicunda* (Fig. 7H) is widespread and very polymorphic (Regalado, 1990, 1995), and it currently includes eight heterotypic synonyms (Kartonegoro, 2022), indicating a potential for further expansion of the species complex or, alternatively, the necessity for a more detailed examination that may lead to a reevaluation of species limits. A study specifically focused on this species complex would be crucial in clarifying the relationships and boundaries within it.

### Medinilla erythrophylla-alliance

Three samples from Myanmar form a clade with full nuclear support (Fig. 4, left). Although only two out of three samples were included in the plastid tree, they also form a well-supported clade (Fig. 4, right). This clade is part of the Western Superclade, spanning from Malesia (primarily Sundaland) to the Afrotropical realm. Stamens are somewhat unequal in these species. *Medinilla pauciflora* (not sampled) shares similarities with *M*. *himalayana*, but it has a more condensed inflorescence and also lacks a dorsal appendage (Clarke, 1879). Both *M*. *himalayana* (Clarke, 1879) and *M*. *griffithii* (Fig. 7I) lack a dorsal spur on the anther, distinguishing them from *M. erythrophylla* which has more typical anthers and is known to have swollen roots. Clausing (1999) placed *M. erythrophylla* in the *M. suberosa*-alliance that included *M. palawanensis*. *Medinilla palawanensis* is very similar in habit to *M. hainanensis* Merr. & Chun (a synonym of *M. erythrophylla*) and *M. ultramaficola* (which has swollen roots). Targeting these taxa is crucial for a better understanding of this clade.

### Medinilla cuneata-alliance

*Medinilla cuneata*, from Sri Lanka, is found in a distinct lineage of its own, sister to the Afrotropical taxa in both nuclear and plastid trees (Fig. 4). Its stems are rather succulent and inflorescences are reduced to single flowers on leafless nodes. Anthers in this species are broadly lanceolate, possessing both ventral lobes and a dorsal spur. *Medinilla anamalaiana* (Western Ghats) and *M. maculata* (Sri Lanka) are morphologically similar species from the same region and probably belong to this clade.

### Medinilla sedifolia-alliance

*Medinilla sedifolia* (Fig. 8B), originating from Madagascar, is resolved as sister to the rest of the Afrotropical *Medinilla* in both nuclear and plastid trees (Fig. 4). Perrier de la Bâthie (1951) and Clausing (1999) treated it in its own subgroup of *M.* sect. *Septatae* and Group 2, respectively. The species strongly resembles *M. beddomei* of the Western Ghats, initially classified as *Triplectrum*. Shared characteristics between these species include a creeping habit, narrow stems, flaky reddish bark, rusty to powdery pubescence on young parts, equal, succulent orbicular leaves, solitary axillary flowers, dimorphic stamens and long blunt ventral and dorsal anther appendages. Another creeping species in this region is *M. prostrata*, with fairly succulent, round leaves, a hairy vestiture, solitary flowers in leaf axils, shorter lobed anther appendages and ovary wholly adherent to the hypanthium (vs. separate). Perrier (1951) did not consider *M. prostrata* very close to *M. sedifolia* because of its distinct anthers and ovary adherence, and he instead placed *M. prostrata* in sect. *Adhaerentes*. Nevertheless, the usefulness of concrescence for phylogenetic inference has not found molecular support (see the *M. viscoides*-alliance discussion below). The connections between these species (plus the pseudotubular-flowered species discussed in the *M. viscoides*-alliance) suggest multiple Madagascar-India connections, warranting further study.

**Figure 8.**
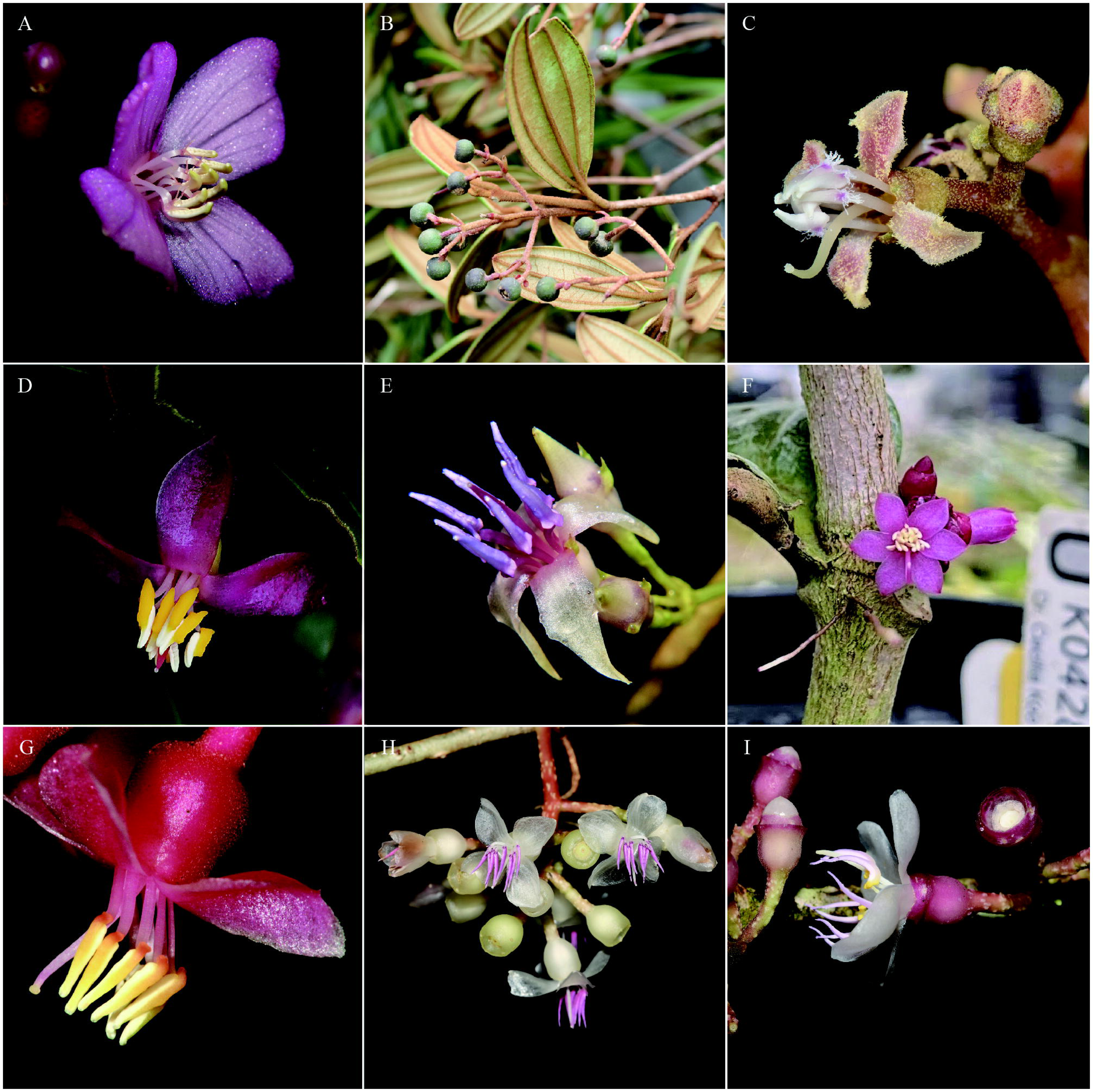
Representatives from major lineages identified within *Medinilla*, continued: (A) *M. fuchsioides*, alliance unknown (Photo: Bathiya Gopallawa, Sri Lanka); (B) *M. sedifolia* from the *M. sedifolia*-alliance (Photo: Marie Selby Botanical Garden, originally from Madagascar); (C) *M. cordifolia* from the *M. viscoides*-alliance (Photo: Maxim Nuraliev, Madagascar); (D) *M. rhodorhachis* from the *M. arfakensis*-alliance (Photo: Shelley James 1867, Solomon Islands); (E) *M. heteromorphophylla* from the *M. anisophylla*-alliance (Photo: Porter Lowry 6861, Vanuatu); (F–I) *M. medinilliana*-alliance; (F) *M. stephanostegia* (Photo: Darin Penneys 2451, Borneo); (G) *M. magnifica* (Photo: Peter Quakenbush, Philippines); (H) *M. quadrifolia* (= *M. trianae*; Photo: Peter Quakenbush 44, Philippines); (I) *M. venosa* (Photo: Peter Quakenbush 61, Philippines).

### Medinilla viscoides-alliance

All species sampled from the Afrotropical realm form a fully supported clade (Fig. 4). *Medinilla mannii* and *M. engleri*, the only two species found on the African continent, are nested among the Malagasy species, indicating their likely origin. *Medinilla sedifolia* discussed above, is sister to all other samples. Excluding *M. sedifolia,* the stamens of the remaining species are easily characterized by long, subulate ventral appendages and a prominent dorsal appendage, exemplified by *M. cordifolia* (Fig. 8C). However, some species from Madagascar, such as *M. papillosa*, exhibit considerable differences. In these species, flowers are nectariferous and relatively large, and the corollas never open fully, forming a pseudotube. The stamens have exceptionally long, broad filaments and much shorter anthers, nearly or entirely lacking appendages. Some species from the Comoro Islands (e.g., *M. fasciculata*), the Western Ghats (e.g., *M. malabarica*) and Sri Lanka (*M. fuchsioides*; Fig. 8A) share this distinctive morphology. The similarities among these disjunct pseudotubular species may be the result of convergent evolution or dispersal across the Indian Ocean. More sampling is needed to verify the placement and relationships of these species.

Swollen/tuberous roots are a noteworthy trait common in this alliance (e.g., *Medinilla mannii*, *M. baronii*, and *M. lophoclada*). They would certainly increase drought tolerance but can be formicarial as well (pers. obs.). Such roots are also observed in *Pachycentria constricta*, *M. ramiflora*, *M. maidenii*, *M. erythrophylla*, and *M. ultramificola*. Thus, it appears to be a widespread trait among the Early Diverging Clades and the Western Superclade.

The first species described in the Afrotropical realm, *Medinilla viscoides*, was classified as *Diplogenea*, as was *M. mannii* (with uncertainty). Both Perrier (1951) and Clausing (1999) grouped *M. viscoides* with *M. chermezonii.* While *M. viscoides* was not sampled in our study, *M. mannii* and multiple samples of *M. chermezonii* were. Both were resolved in this clade, providing strong molecular evidence for including *Diplogenea* in *Medinilla*. Species in this clade also correspond to two sections, *M.* sect. *Septatae* and *M.* sect. *Adhaerentes*. These were based on the degree of adherence of the hypanthium to the ovary (Perrier, 1951). Both sections are represented by multiple taxa in our study, and they do not receive molecular support. Additionally, species in this clade correspond to Clausing’s (1999) Group 2, which includes four alliances: *M. sedifolia*, *M. parvifolia*, *M. ericarum*, and *M. homblotii*. The *M. sedifolia*-alliance is discussed above, the *M. parvifolia*-alliance does not gain molecular support, and sampling was insufficient to test the *M. homblotii* and *M. ericarum* alliances. Weak branch support, particularly in the plastid phylogeny, combined with short branch lengths (Supplementary Fig. S3 & S4) and discordance among the phylogenies, prevents further division of this group. Therefore, increased sampling, detailed morphological comparisons, and a comprehensive revision of species are critically needed. Additionally, incorporating more plastid loci, such as the entire plastome, is essential for a deeper understanding of the internal relationships within this group.

### Medinilla arfakensis-alliance

Three samples from New Guinea, the Bismarck Archipelago, and the Solomon Islands (Fig. 8D) were resolved in a clade fully supported by both nuclear and plastid trees (Fig. 5). However, species identification is challenging due to the limited knowledge of taxa in this region. A comprehensive revision and more fieldwork are urgently needed. The clade is sister to the *Medinilla anisophylla*-alliance discussed below. Both of these are sister to the *M*. *medinilliana*-alliance, and collectively form the Eastern Superclade. Regalado (1990) previously considered many of the species in New Guinea as part of his *M. magnifica*-alliance. Clausing (1999) later expanded the *M. magnifica*-alliance, including several species from New Guinea (e.g., *M. arfakensis*). However, this expanded alliance was found to be polyphyletic with sampled species split between the *M*. *arfakensis* and *M*. *medinilliana* alliances. The *M*. *arfakensis*-alliance tends to have a prominent dorsal appendage and small or absent ventral appendages, while similar species in the *M*. *medinilliana*-alliances tend to have a small dorsal spur and more prominent ventral lobes. Undoubtedly, there are more species from New Guinea and surrounding islands belonging to this clade. To gain a better understanding of this group, more extensive sampling and foundational taxonomic work are essential.

### Medinilla anisophylla-alliance

A clade consisting of seven taxa from the Solomon Islands is fully supported by both phylogenies and is sister to the *Medinilla arfakensis*-alliance (Fig. 5). These species are characterized by being hairy climbers with strongly anisophyllous leaves, prominent and persistent floral bracts, hairy hypanthia, and robust dorsal appendages on the anthers (e.g., Fig. 8E). When present, the ventral appendages are much less prominent than the dorsal appendage. Despite differences in anther morphology, these taxa share strong similarities with some species resolved in or associated with the *M*. *medinilliana*-alliance (see discussion below). For example, *M. cephalantha* was compared to *Cephalomedinilla* (Merrill & Perry, 1943). Another species from Vanuatu, undoubtedly belonging in this clade, is *M. heterophylla*, which was likened to *Dactyliota* (Gray, 1854). These apparent similarities are likely the result of convergent evolution. Additional taxa from the Bismarck Archipelago (e.g., *M. pubiflora* Merr. & L.M.Perry; NGF 31506, BISH & L) and the Solomon Islands (e.g., *M. sessilis*) likely belong to this clade. Similarly, taxa from Micronesia (i.e., Kosrae), Fiji, Wallis and Futuna, Samoa, and American Samoa are likely part of this clade. Notably, *M. calliantha* was not resolved as monophyletic and might need revision.

### Medinilla medinilliana-alliance

A large clade primarily composed of Philippine taxa, including the type species (*Medinilla medinilliana*), was resolved with full support by both datasets (Figs. 5 & 8F–I). This clade extends its distribution to include Vietnam, Borneo, Guam, New Guinea, the Solomon Islands, and Australia, making this the most widespread and species-rich clade identified in this study. The alliance is characterized by an exceptionally high degree of polymorphism, rendering it challenging to precisely define. The considerable diversity within this clade is evident from the various genera that were historically associated with it. As earlier noted in the *Medinilla* overview, species once classified as *Hypenanthe* (*M. venosa*), *Dactyliota* (*M. setigera*), and *Carionia* (*M. whitfordii*) were resolved within this alliance. Regalado (1995) included the first two in his Group 10, where he also placed *Cephalomedinilla* (not tested). Morphological characteristics, including habit, stem, leaf, inflorescence, and floral details are consistent with this group. Therefore, it is reasonable to include *Cephalomedinilla* within this alliance as well. Many species from other described groups were resolved within this clade, especially those from Regalado (1990, 1995) and Clausing (1999; Group 1). Only the small *M. setphanostegia*-alliance (Fig. 8F; _J species sampled) found support without the need for modification.

In general, the anthers of taxa within the *Medinilla medinilliana*-alliance are narrow and curved, consisting of two distinct colors (e.g., yellow and purplish). The ventral lobes tend to be more prominent than the dorsal spur or both ventral and dorsal appendages are long and conspicuous. Although, anthers in the *M. rubicunda*, *M. erythrophylla*, *M. cuneata*, and *M. sedifolia* alliances of the Western Superclade may share some similarities, species in the *M*. *medinilliana*-alliance can be distinguished by additional traits. These distinctive features include setose nodes, inflorescences with regularly whorled branches, conspicuous bracts, membranous calyx rim, and/or 6-merous flowers. Whorled leaves, a hairy or thick hypanthium, heteranthery, and/or bicolored anthers serve as reliable indicators of this clade as well, although these traits may occasionally appear in the Western Superclade.

Despite the widespread phylogenetic discordance observed between the two phylogenies, some clades within the nuclear tree exhibit morphological coherence. For instance, the *M. medinilliana*-*M. whitfordii* clade, supported by the nuclear tree (NAT: LPP 0.64), consists of terrestrial shrubs or climbers with a thick hypanthium (e.g., > 1 mm). Many also have whorled leaves, 5 or 6-merous flowers, and heteranthery. Another coherent clade, the *M. erythrotricha*-*M. disparifolia* clade (NAT: LPP 1), are all terrestrial shrubs or climbers as well. Bracts tend to be prominent, and the hypanthium tends to be hairy with a membranous calyx rim. The *M. magnifica*-*M. clementis* and *M. teysmannii*-*M. theresae* clades tend to be epiphytic shrubs with setose nodes and many-flowered inflorescences. However, additional targeted systematic work is needed to further characterize and understand these groups. Several species, including *M. pendula*, *M. multiflora*, *M. setigera*, and members of the *M. quadrifolia*-*M. Medilliana* clade are in need of taxonomic review due to the presence of synonyms and wide distribution in the Philippines and beyond.

## TAXONOMIC TREATMENT

In this section, *Medinilla* is redefined. Synonymous genera are listed, including *Pachycentria*, *Plethiandra*, their synonyms, and type. A new diagnosis, new description, distribution, and notes on the correct authorship of *Medinilla* are given. To provide *Medinilla* names for *Pachycentria* taxa, eight new name combinations are effected and one name is reinstated. To provide *Medinilla* names for *Plethiandra* species, four new names are provided, three new combinations are effected, and one name is reinstated. Accepted species follow Clausing (2000) and Kadereit (2005).

***Medinilla*** Gaudich. ex DC., Prodr. [A. P. de Candolle] 3: 167. 1828 – Type: *Melastoma medinilliana* Gaudich. (= *Medinilla medinilliana* (Gaudich.) Fosberg & Sachet).
= *Diplogenea* Lindl., Quart. J. Sci. Lit. Arts 1828(2): 122. 1828 – Type: *Diplogenea viscoides* Lindl. (= *Medinilla viscoides* (Lindl.) Triana).
= *Pachycentria* Blume, Flora 14: 519. 1831, **syn. nov.** – Type: *Pachycentria constricta* (Blume) Blume, Flora 14: 520. 1831 (= *Medinilla constricta* (Blume) Quakenbush & Luo Chen, this paper).
= *Pogonanthera* Blume, Flora 14: 520. 1831, **syn. nov.** – Type: *Pogonanthera pulverulenta* (Jack) Blume (= *Medinilla pulverulenta* (Jack) Quakenbush & Luo Chen, this paper).
*= Triplectrum* D.Don ex Wight & Arn., Prodr. Fl. Ind. Orient. 1: 324. 1834 – Type: *Triplectrum radicans* D.Don ex Wight & Arn. (= *Medinilla beddomei* C.B.Clarke).
= *Dactyliota* Blume, Mus. Bot. 1(2): 21. 1849 – Type: *Dactyliota bracteata* (Blume) Blume (= *Medinilla bracteata* (Blume) Blume).
= *Hypenanthe* Blume, Mus. Bot. 1(2): 21. 1849 – Type: *Hypenanthe venosum* (Blume) Blume (= *Medinilla venosa* Blume).
= *Carionia* Naudin, Ann. Sci. Nat., Bot. sér. 3, 15: 311, t. 15. 1851 – Type: *Carionia elegans* Naudin (= *Medinilla coronata* Regalado).

= *Erpetina* Naudin, Ann. Sci. Nat., Bot. sér. 3, 15: 299. 1851 – Type: *Erpetina radicans* Naudin (= *Medinilla erpetina* (Naudin) Triana).
= *Plethiandra* Hook.f., Gen. Pl. [Bentham & Hooker f.] 1(3): 772. 1867, **syn. nov.** – Type: *Plethiandra motleyi* Hook.f. (= *Medinilla polystamina* Quakenbush, this paper).
= *Medinillopsis* Cogn., Monogr. Phan. [A.DC. & C.DC.] 7: 603. 1891, **syn. nov.** – Type (designated by Kadereit in Edinburgh J. Bot. 62(3): 131. 2005): *Medinillopsis beccariana* Cogn. (= *Medinilla incognita* Quakenbush, this paper).
= Cephalomedinilla Merr., Philipp. J. Sci., C 5: 204. 1910 – Type: Cephalomedinilla anisophylla Merr. (= Medinilla microcephala Regalado).
– “Medinilla Gaudich.”, isonym, in Gaudichaud-Beaupré, Voy. Uranie, Bot. pt. 11: plate 106. 1829; pt. 12: 484. 1830.

Diagnosis: *Medinilla* can be distinguished from all other Sonerileae by the combination of typical xylem (vs. lobed in cross-section) and berry fruit. *Medinilla* can be distinguished from the Dissochaeteae by its wood anatomy (e.g., distinctly scalariform intervessel pits; van Vliet, 1981), the absence of interpetiolar ridges (Veldkamp, 1978), and leaf venation (generally suprabasal acrodromous vs. basal acrodromous). At present, *Medinilla* can be phylogenetically defined as the most exclusive clade containing *M. medinilliana* and *M. nubicola*.

Description: Terrestrial shrubs/small trees, lianas, primary hemiepiphytes, and epiphytes; roots sometimes swollen; stems terete or tetragonal, sometimes 4–8-winged, glabrous or pubescent; nodes often thickened, with or without setae; leaves opposite or whorled, glabrous or pubescent, sessile or petiolate, sometimes with pseudostipules, strongly anisophyllous to equal, usually coriaceous or fleshy, venation generally suprabasal acrodromous, with 1–many nerves, base variable (e.g., peltate, auriculate, obtuse, acute), apex variable (e.g., retuse, obtuse, acute), margin entire; inflorescences terminal, axillary, or cauline, cymose, 1–many-flowered, diffuse to densely congested, solitary to fascicled, lax to erect, with or without showy bracts and bracteoles, glabrous or pubescent; flowers 4–6(–7)-merous; hypanthium variable (e.g., ovoid, campanulate, cylindrical, urceolate), occasionally bumpy or with long outgrowths, glabrous or pubescent; calyx rim (limb) variable (e.g., truncate, dentate, regularly or irregularly lobed, variously flared); petals broadly or narrowly oblique, apex rounded or pointed, white, pink, lavender, orange, or red, recurved, spreading, cupped, or pseudotubular; stamens generally double the petal number, e.g. 8–12, but more than double (polyandrous) in the *Plethiandra* clade; filaments strap-shaped, pale; anthers white, yellow, pink, red, blue, purple, or a combination thereof, isomorphic, subequal, or dimorphic, opening by 1(–2) pores, variously arranged (e.g., evenly distributed, in one or two groups); pedoconnective not or hardly produced at the base, with or without dorsal and ventral appendages; dorsal appendage forming a triangular plectrum, subulate or spatulate spur of various lengths, split into two, frayed, or presenting as a tuft of hairs; ventral appendages long or short, subulate or lobed; ovary partially to wholly adnate to the calyx, usually separated by extra-ovarian chambers (corresponding to stamen number and length of anther/filament), 4–6-locular, placentation axial, apex concave or convex, glabrous; style straight or hooked; stigma punctate to capitate; fruit baccate (an accessory fruit with a fleshy hypanthium), ovoid, globose, urceolate, or ellipsoid, green, white, pink, yellow, orange, red, or some combination thereof when immature, green, blue, dark purplish-black when mature; seeds few to many, minute to ∼1.5mm, semi-ovate to irregularly ovoid or angular, testa smooth to papilate, testa cells interdigitate to comb-shaped, hilum basal, raphe often evident.

Distribution: Afrotropical, Indomalayan, Australasian, and Oceanian biogeographic realms, i.e., throughout much of the wet Paleotropics.

Notes: Authorship for *Medinilla* is often erroneously cited. For example, Regalado (1990, 1995) cited Gaudich. (1826), Veranso-Libalah et. al (2022) cited Gaudich. (1830), and Kartonegoro (2023) cited Gaudich. (1828). However, Augustine Pyramus de Candolle was the first to validly publish *Medinilla* in mid-March 1828. De Candolle had early access to Gaudichaud-Beaupré’s material, which was not published until 1829 and 1830 (see references in list of synonyms above). De Candolle cited Gaudichaud-Beaupré’s unpublished work, and provided his own, at times conflicting, description. Thus, the correct authorship is Gaudich. ex DC, and *Medinilla* Gaudich. is an isonym (Art. 6, Note 2). For a detailed explanation of the issue, see Bodegom and Veldkamp (2001), which we verified with Kanchi Gandhi (pers. comm.), Senior Nomenclature Registrar and Bibliographer at Harvard University. *Diplogenea* was published soon after, in October of 1828 and does not have priority.

List 1. *Medinilla* names for accepted *Pachycentria* taxa (refer to Clausing [2000] for full list of heterotypic synonyms)

***Medinilla constricta*** (Blume) Quakenbush & Luo Chen, **comb. nov.** ≡ *Melastoma constrictum* Blume, Bijdr. Fl. Ned. Ind. 17: 1072. 1826 ≡ *Pachycentria constricta* (Blume) Blume, Flora 14: 520. 1831. – Type: Indonesia, Java, *Blume s.n.* (lectotype L barcode L908.132-896, designated by Clausing in Blumea 45(2): 351. 2000).
***Medinilla glauca*** subsp. ***glauca*** (Triana) Quakenbush & Luo Chen, **comb. nov.** ≡ *Pachycentria glauca* Triana, Trans. Linn. Soc. London 28(1): 89. 1871 – Type: Malaysia, Sarawak, *Beccari 415* (holotype FI; isotype K).
***Medinilla glauca*** subsp. ***maingayi*** (C.B.Clarke) Quakenbush & Luo Chen, **comb. nov.** ≡ *Pachycentria glauca* subsp. *maingayi* (C.B.Clarke) Clausing, Blumea 45(2): 356. 2000 ≡ *Pachycentria maingayi* (C.B.Clarke) J.F.Maxwell, Gard. Bull. Singapore 31(2): 203. 1978 ≡ *Medinilla maingayi* C.B.Clarke, Fl. Brit. India [J. D. Hooker] 2(6): 549. 1879 – Type: Singapore, *Maingay 806 (3329)* (syntype K); Malaysia, Malacca, *Maingay 807 (2960)* (syntype K).
***Medinilla hanseniana*** (Clausing) Quakenbush & G.Kadereit, **comb. nov.** ≡ *Pachycentria hanseniana* Clausing, Blumea 45(2): 356. 2000 – Type: Indonesia, Kalimantan, Tengah, Kualakuayan, *Hansen 1336* (holotype C).
***Medinilla microsperma*** (Becc.) Quakenbush & Luo Chen, **comb. nov.** ≡ *Pachycentria microsperma* Becc., Malesia 2: 238, t. 58, Fig. 1-9. 1886; Cogn. in DC. Monog. Phan. 7: 609. 1891 – Type: Malaysia, Sarawak, *Beccari 404* (holotype FI; isotype K).
***Medinilla microstyla*** (Becc.) Quakenbush & Luo Chen, **comb. nov.** ≡ *Pachycentria microstyla* Becc., Malesia 2: 239. 1886; Cogn. in DC. Monog. Phan. 7: 606. 1891 – Type: Malaysia, Sarawak, near Kuching, *Beccari 604* & *403* (syntypes FI).
***Medinilla pulverulenta*** (Jack) Quakenbush & Luo Chen, **comb. nov.** ≡ *Melastoma pulverulentum* Jack, Trans. Linn. Soc. London 14(1): 19. 1823 ≡ *Pogonanthera pulverulenta* (Jack) Blume, Flora 14: 521. 1831 ≡ *Pachycentria pulverulenta* (Jack) Clausing, Blumea 45(2): 362. 2000 – Type: Sumatra, *Jack s.n.* (holotype presumably lost); Malaysia, Malacca, Cape Rochado, 1822, *Wallich 4086(A)* (neosyntype K barcode K001038107 [image!], **designated here**), *Wallich 4086(B)* (neosyntype K barcode K001038108 [image!], **designated here**).
***Medinilla varingiifolia*** (Blume) Nayar, Blumea 18: 567. 1970 ≡ *Melastoma varingiaefolium* Blume, Bijdr. Fl. Ned. Ind. 17: 1071. 1825 ≡ *Pachycentria varingaefolia* [Blume] Blume, Flora 14: 520. 1831 – Type: Indonesia, Java, *Kuhl & van Hasselt s.n.* (holotype L barcode L908.132-158; isotypes L barcode L908.132-168, L barcode L908.132-178).
***Medinilla vogelkopensis*** (Clausing) G.Kadereit & Luo Chen, **comb. nov.** ≡ *Pachycentria vogelkopensis* Clausing, Blumea 45(2): 367. 2000 – Type: New Guinea, Vogelkop Peninsula, Mt. Nettoti, path Andjai-Wekari at 1650m, *Van Royen & Sleumer 7902* (holotype L; isotype A).

List 2. *Medinilla* names for *Plethiandra* species (refer to Kadereit [2005] for full list of heterotypic synonyms)

***Medinilla hookeri*** (Stapf) Quakenbush & Luo Chen, **comb. nov.** ≡ *Plethiandra hookeri* Stapf, Trans. Linn. Soc. London, Bot. 4(2): 163. 1894 – Type: Malaysia, Sabah, Mt. Kinabalu, Aug. 1892, *Haviland 1169* (lectotype K, designated by Nayar in Reinwardtia 9(1): 147. 1974; isolectotype SAR.
***Medinilla incognita*** (Cogn.) Quakenbush, **nom. nov.** ≡ *Medinillopsis beccariana* Cogn., Monogr. Phan. [A.DC. & C.DC.] 7: 603. 1891 ≡ *Plethiandra beccariana* (Cogn.) Merr., J. Straits Branch Roy. Asiat. Soc. 84(Spec. No.): 448. 1921 – Type: Malaysia, Sarawak, Bintulu, Sep. 1867, *Beccari 4004* (lectotype FI; isolectotype K).
***Medinilla stapfii*** Quakenbush, **nom. nov.** ≡ *Plethiandra sessilis* Stapf, Hooker’s Icon. Pl. 25: t. 2418. 1895 – Type: Malaysia, Sarawak, Penrissen, Jun. 1890, *Haviland 6893* (lectotype K).
***Medinilla migrans*** Quakenbush, **nom. nov.** ≡ *Medinillopsis sessiliflora* Cogn., Monogr. Phan. [A.DC. & C.DC.] 7: 603. 1891 ≡ *Plethiandra sessiliflora* (Cogn.) Merr., J. Straits Branch Roy. Asiat. Soc. 84(Spec. No.): 449. 1921 – Type: Malaya, 1866, *Beccari s.n.* (lectotype FI; isolectotype K).
***Medinilla polystamina*** Quakenbush, **nom. nov.** ≡ *Plethiandra motleyi* Hook.f., Gen. Pl. [Bentham & Hooker f.] 1(3): 772. 1867 – Type: Malaysia, Sabah, Labuan, *Motley 380* (lectotype K).
***Medinilla rejangensis*** (Stapf) Quakenbush & Luo Chen, **comb. nov.** ≡ *Plethiandra rejangensis* Stapf, Hooker’s Icon. Pl. 25: sub t. 2418. 1895 – Type: Malaysia, Sarawak, Rejang, Sibu., *Haviland 545* (lectotype K). = *Plethiandra cuneata* Stapf, Hooker’s Icon. Pl. 25: sub t. 2418. 1895 – Type: Malaysia, Sarawak, Selabat rock, sea coast, Mar. 1891, *Haviland 179* (lectotype K; isolectotypes BM, SAR, SING).
***Medinilla robusta*** Cogn. ≡ *Plethiandra* robusta (Cogn.) Nayar, Reinwardtia 9(1): 148. 1974 – Type: Malaysia, Sarawak, Kuching, *Beccari 542* (lectotype FI; isolectotype K, designated by Kadereit in Edinburgh J. Bot. 62(3): 137. 2005); Malaysia, Sarawak, Kuching, *Beccari 851* (syntype FI); Malaysia, Sarawak, Bintulu, Nov. 1867, *Beccari 4049* (syntype FI).
***Medinilla tomentosa*** (G.Kadereit) G.Kadereit, **comb. nov.** ≡ *Plethiandra tomentosa* G.Kadereit, Edinburgh J. Bot. 62(3): 141. 2005 – Type: Malaysia, Sarawak, Lambir National Park, 1983, *B. Lee S.46581* (holotype AAU; isotype KEP).

Notes: *Medinilla incognita* is a replacement name for *Medinillopsis beccariana*, because *Medinilla beccariana* Cogn. already exists for another species. *“Incognita*” is in reference to this species’ long-hidden identity as *Medinilla*. *Medinilla stapfii* is a replacement name for *P. sessilis*, because *M. sessilis* Merr. & L.M.Perry already exists. It is named in recognition of Otto Stapf, the original author of this species and in recognition of his contributions to the clade. *Medinilla migrans* is a replacement name for *P. sessiliflora*, because *M. sessiliflora* Regalado already exists. “*Migrans*” recognizes that this species is the only *Plethiandra* species found outside of Borneo. *Medinilla polystamina* is a replacement name for *P. motleyi*, because *M. motleyi* Hook.f. ex Triana already exists. The epithet from the heterotypic synonym *P. acuminata* Merr. cannot be used, because *M. acuminata* Merr. already exists. *Medinilla polystamina* is the type species of *Plethiandra*, and the name acknowledges the most unique feature of this group–its many stamens. *Medinilla robusta* is reinstated as the accepted name for *P. robusta*. *Medinilla rejangensis* is the new name combination for *P. cuneata*; because *M. cuneata* (Thwaites) K.Bremer & Lundin already exists, and *P. rejangensis* Stapf is the next, earliest, legitimate name for this taxon.

## CONCLUSIONS

We provide a substantial advancement towards a well-sampled and resolved phylogeny of fleshy-fruited Sonerileae. The *Medinilla*-alliance, characterized by a typical wood stele and soft, juicy berries, is monophyletic. However, *Medinilla* in its current circumscription is paraphyletic because *Pachycentria* and *Plethiandra* are nested within the clade. A number of taxonomic changes are proposed and outlined in the taxonomic treatment section. Including these *Pachycentria* and *Plethiandra*, 15 major lineages within *Medinilla* are identified and lay the basic structure for a comprehensive infrageneric classification system. More sampling, especially from Madagascar, India, Sundaland, and New Guinea is needed to explore the limits and internal relationships of these lineages further.

In contrast to the typical wood stele of the *Medinilla*-alliance, the *Heteroblemma*-alliance, characterized by lobed stele and various fruit types, is polyphyletic. *Kendrickia* is sister to an Afrotropical superclade, while *Heteroblemma* and *Catanthera* belong to a mostly Asian superclade. Plastid sequences of *Kendrickia* are still lacking and will help verify the relationship. Both *Heteroblemma* and *Catanthera* could benefit from greater sampling, especially from east of the Wallace Line, which has no representation yet. *Heteroblemma* in particular requires further attention, because it was only resolved as monophyletic in the nuclear tree. Plastid sequences showed Bornean species with a closer relationship to *Phyllagathis* species. The origin of this discordance needs further exploration.

## AUTHOR CONTRIBUTIONS

JPQ, LC, MCVL, and GK designed the project. JPQ and LC performed the research. LC and JPQ conducted data collection, analysis, and interpretation, with assistance from MCVL, GK, DSP, YL, and DY. JPQ and LC drafted the manuscript. All authors contributed to the revision process.

## ACKNOWLEDGEMENTS

The authors would like to thank MJG, MO, NY, K, CSH, E, TNM, the Makiling Center for Mountain Ecosystems, Marie Selby Botanical Gardens, Hortus Botanicus Leiden, Munich Botanical Garden, NSF-DEB award numbers 1754697, 1754667, 0950207, 1457702, Nina Peck, Jiro Adorador, and Maxim Nuraliev for providing samples; Martina Silber for aiding in DNA extraction. This project is funded by the German Science Foundation (DFG), project number KA1816/12-1 and VE 1291/1-1. JPQ would also like to thank The Garden Club of America, Western Michigan University, the Gwen Frostic Doctoral Fellowship, Kalamazoo Garden Club, and International Association for Plant Taxonomy for funding. This article is part of the PhD thesis of JPQ and LC.

## DATA AVAILABILITY STATEMENT

All sequencing data generated in this study are deposited in the National Center for Biotechnology Information (NCBI) Sequence Read Archive, and the phylogenetic trees and alignments are deposited in Zenodo.

## SUPPORTING INFORMATION

Supplementary Table S1. Detailed voucher information and accession numbers for SRA and ENA.

Supplementary Table S2. Gene recovery statistics for all samples.

Supplementary Fig. S3. DISCO-ASTRAL phylogram. Local Posterior Probabilities are shown below the branches.

Supplementary Fig. S4. Maximum Likelihood plastid phylogram. Bootstrap support values are shown below the branches.

