## Supplementary figures and images for "Systematics of the fleshy-fruited Sonerileae (Melastomataceae)"

### Supplementary Fig. S3

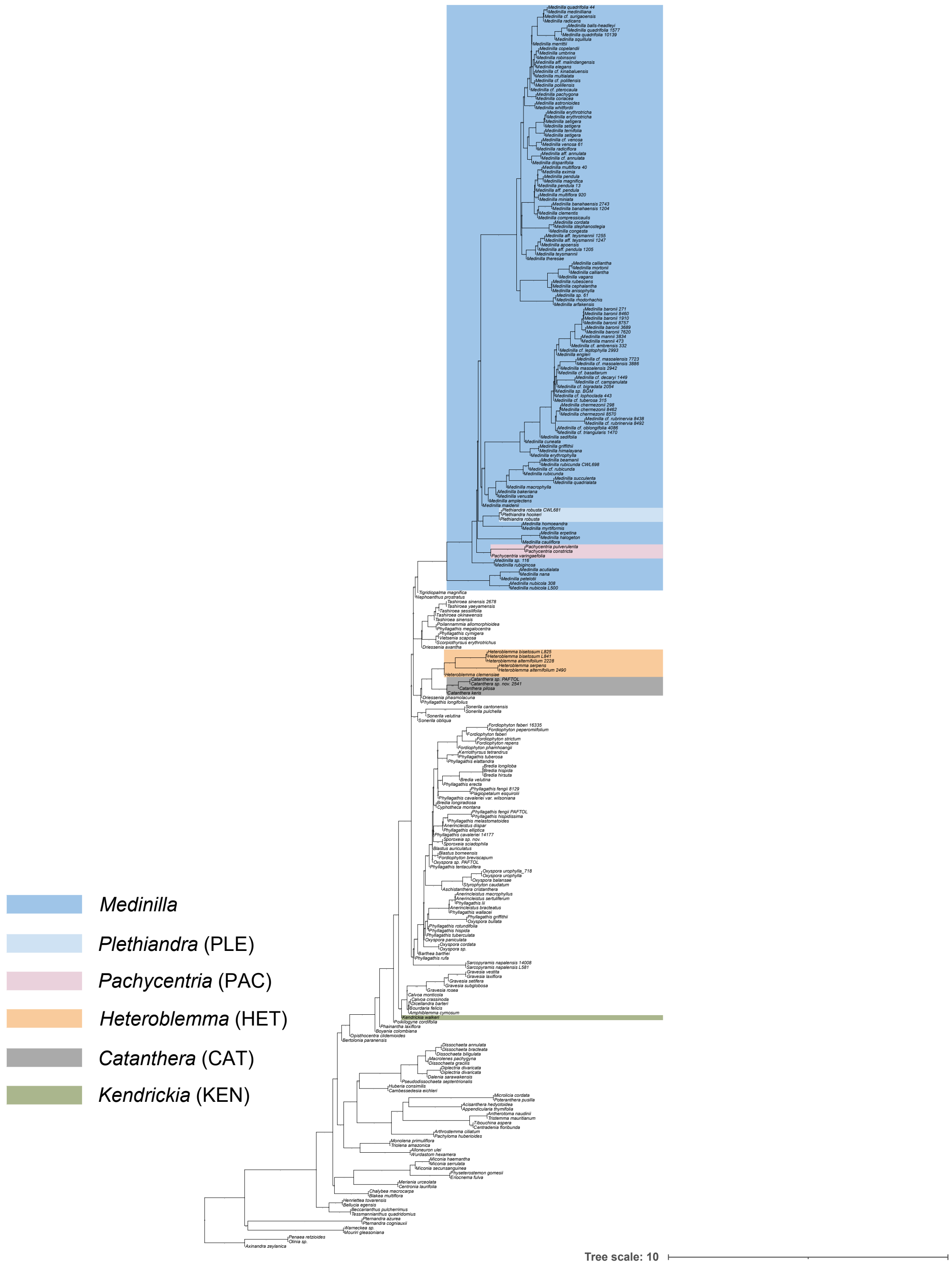

### Supplementary Fig. S4

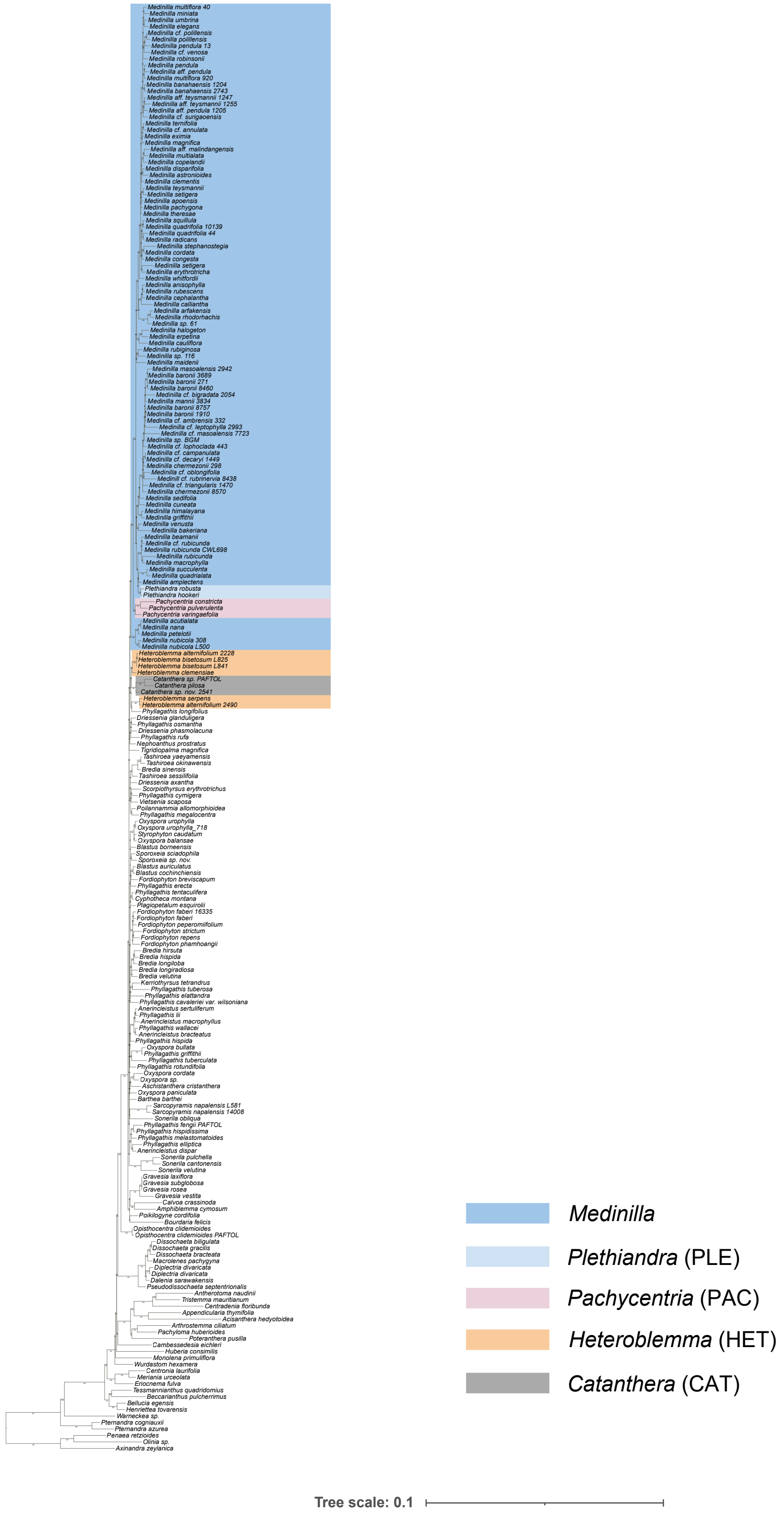
